# Genome-wide Analysis Reveals Novel Regulators of Growth in *Drosophila melanogaster*

**DOI:** 10.1101/017855

**Authors:** Sibylle Chantal Vonesch, David Lamparter, Trudy FC Mackay, Sven Bergmann, Ernst Hafen

**Author notes:** Current Address: Genome Biology Unit, European Molecular Biology Laboratories, Heidelberg, Germany (Sibylle Chantal Vonesch). Corresponding author (EH).

## Abstract

Organismal size depends on the interplay between genetic and environmental factors. Genome-wide association (GWA) analyses in humans have implied many genes in the control of height but suffer from the inability to control the environment. Genetic analyses in *Drosophila* have identified conserved signaling pathways controlling size; however, how these pathways control phenotypic diversity is unclear. We performed GWA of size traits using the *Drosophila* Genetic Reference Panel of inbred, sequenced lines. We find that the top associated variants differ between traits and sexes; do not map to canonical growth pathway genes, but can be linked to these by epistasis analysis; and are enriched for genes and putative enhancers. Performing GWA on well-studied developmental traits under controlled conditions expands our understanding of developmental processes underlying phenotypic diversity.

**AUTHOR SUMMARY:** Genetic studies in *Drosophila* have elucidated conserved signaling pathways and environmental factors that together control organismal size. In humans, hundreds of genes are associated with height variation, but these associations have not been performed in a controlled environment. As a result we are still lacking an understanding of the mechanisms creating size variability within a species. Here, under carefully controlled environmental conditions, we identify naturally occurring genetic variants that are associated with size diversity in *Drosophila*. We identify a cluster of associations close to the *kek1* locus, a well-characterized growth regulator, but otherwise find that most variants are located in or close to genes that do not belong to the conserved pathways but may interact with these in a biological network. Many of these genes have a conserved role in humans. We validate 33 novel growth regulatory genes that participate in diverse cellular processes, most notably cellular metabolism and cell polarity. This study is the first genome-wide association analysis of natural variants underlying size in *Drosophila* and our results complement the knowledge we have accumulated on this trait from mutational studies of single genes.

## INTRODUCTION

How animals control and coordinate growth among tissues is a fundamental question in developmental biology. A detailed mechanistic but global understanding of the processes taking place during normal physiological development is furthermore relevant for understanding pathological growth in cancers. Classical genetic studies in *Drosophila* have revealed core molecular mechanisms governing growth control and have shed light on the role of humoral factors and the environment on adult size (1,2,3,4). Two major pathways regulate size, the Insulin/TOR pathway, which couples systemic growth to nutrient availability; and the Hippo tumor suppressor pathway, which controls cell survival and proliferation in developing organs (5,6,7). However, growth control is complex (8,9,10), and the interactions between components of these pathways with each other and with unknown molecules and extrinsic factors remain poorly understood. Studies focusing on single or a few genes can only capture individual aspects of the entire system of networks underlying this trait, which is especially problematic when individual alleles have subtle and context-dependent effects (11,12). Therefore, global genome-wide approaches are needed for a better understanding of the genetic control of size. One genome-wide approach is to study multifactorial natural genetic perturbations as they occur in a segregating, phenotypically diverse population.

Artificial selection experiments have revealed that naturally occurring populations of *Drosophila melanogaster* show abundant genetic variation for size, with heritabilities approaching 50% (13). Usually selection for size results in correlated responses in the same direction for all body parts and overall weight, indicating a common genetic architecture (14). Selection responses differ between populations but not between sexes (15). Body size is an important component of fitness in *D. melanogaster* since there are parallel clines in body size and correlated traits clines across different continents (17,18). Loci on chromosome *3R* and *2R* are, respectively, associated with body size and wing area (17, 18); interestingly, the majority of the *3R* loci seem to be located within the polymorphic chromosomal inversion *In(3R)Payne* (19, 20). Candidate genes and variants associated with size within *In(3R)Payne* include *hsr-omega*, the microsatellite loci DMU25686 and AC008193, and genes in the Insulin signaling pathway (*InR*, *Tsc1*, *Akt1*) (21, 22). Similarly, the frequency of the polymorphic inversion *In(2L)t* is associated with a body size cline across several continents; genes in the IIS/TOR pathway (*chico*, *Pten*, *Tor*) are located in the inversion region and *Pi3K21B* and *Idgfs 1-3* are located immediately proximal to it (22). Naturally segregating alleles in *smp-30* (*Dca*) and *InR* have been causally associated with body weight (23, 24).

Recently, a long term selection experiment identified hundreds of loci with allele frequency differences between large and small populations (16), indicating that the genetic basis of naturally occurring variation in size is highly polygenic. Candidate loci were enriched for genes implicated in post-embryonic development, metamorphosis and cell morphogenesis. The genes included components of the EGFR, Hippo and many other growth pathways, as well as canonical IIS/TOR signaling genes. Therefore, dissecting the genetic basis of naturally occurring variation in body size has the potential to uncover novel variants in known loci affecting body size as well as identify novel genes.

The advent of next-generation sequencing technology has enabled the rapid and relatively cheap acquisition of complete genome sequences, and thereby the generation of very dense genotype information that enables genome-wide association (GWA) mapping with a much higher resolution than previously possible. GWA studies aim to link variation in quantitative traits to underlying genetic loci in populations of unrelated individuals genome-wide (25,26). GWAS have been pioneered (27,28) and widely applied in humans and are now a routinely used tool in model organisms such as *Arabidopsis* (29,30), *Drosophila* (31,32,33) and mouse (34) as well as in various crop (35,36) and domestic animal species (37,38,39,40,41), where they have substantially broadened our understanding of the genetics of complex traits. To date there are no GWA analyses of size in *Drosophila*, but GWA studies of height have revealed that many loci with small effect contribute to size variation in human populations (9,10,42,43), which contrasts with a much simpler genetic architecture of size in domestic animals, where as a consequence of breeding few loci have relatively large effect sizes that jointly explain a large proportion of size variation (38,44). Although many loci affecting human height have been identified by GWA analyses, deducing the underlying molecular mechanisms by which they affect size is challenging. Larger genome regions and not single genes are mapped; uncontrolled environmental variability makes it difficult to identify causal links between genotype and phenotype; and functional validation cannot be performed in humans (12,28,45,46,47).

In contrast to human studies, GWA studies in model organisms benefit from the feasibility of functional validation, more stringent environmental control and, when using inbred strains, the possibility of measuring many genetically identical individuals to obtain an accurate estimate of the phenotype for a given trait and genotype. All three factors can substantially improve the power of a GWA analysis. The establishment of the inbred, sequenced lines of the *Drosophila* Genetic Reference Panel (DGRP) (48,49) has made GWA analysis in *Drosophila* widely applicable. The DGRP lines harbor the substantial natural genetic variation present in the original wild population and show copious phenotypic variation for all traits assayed to date (31,32,33,48,50,51).

Here, we used the DGRP to perform single-and two-locus associations for size-related developmental traits in *Drosophila.* We find pervasive trait and sex-specificity of top variants, validate a substantial number of novel growth regulators, and extend our knowledge of the genetic control of size beyond existing growth regulatory networks.

## RESULTS

### Quantitative genetic analysis of size

We cultured 143 DGRP lines under conditions we had previously shown to reduce environmental influences on size (S1 Table, Fig. 1a) and measured five body and 21 wing traits (S1 Table, Fig. 1b). The cross trait genetic correlations were positive and generally high among all features except small veins and areas that were difficult to quantify accurately, indicating shared genetic architecture of the various size measures. We observed two modules of higher correlation, one formed by wing traits and the second by head/thorax traits (Fig. 1c, S1 Fig.), indicating that the genetic architecture is more similar among wing features and among head/thorax features than between traits of the wing and head/thorax. Principal component analysis (PCA) of 23 of the 26 size traits (L1, L6 and iarea8 were excluded since measuring these traits accurately was very difficult) revealed that the first two PCs explained nearly 75% of the observed phenotypic variation. The first component reflected an overall size element and the second component separated wing from head/thorax traits (Fig. 1d-f, S1 Fig.).

**Figure 1.**
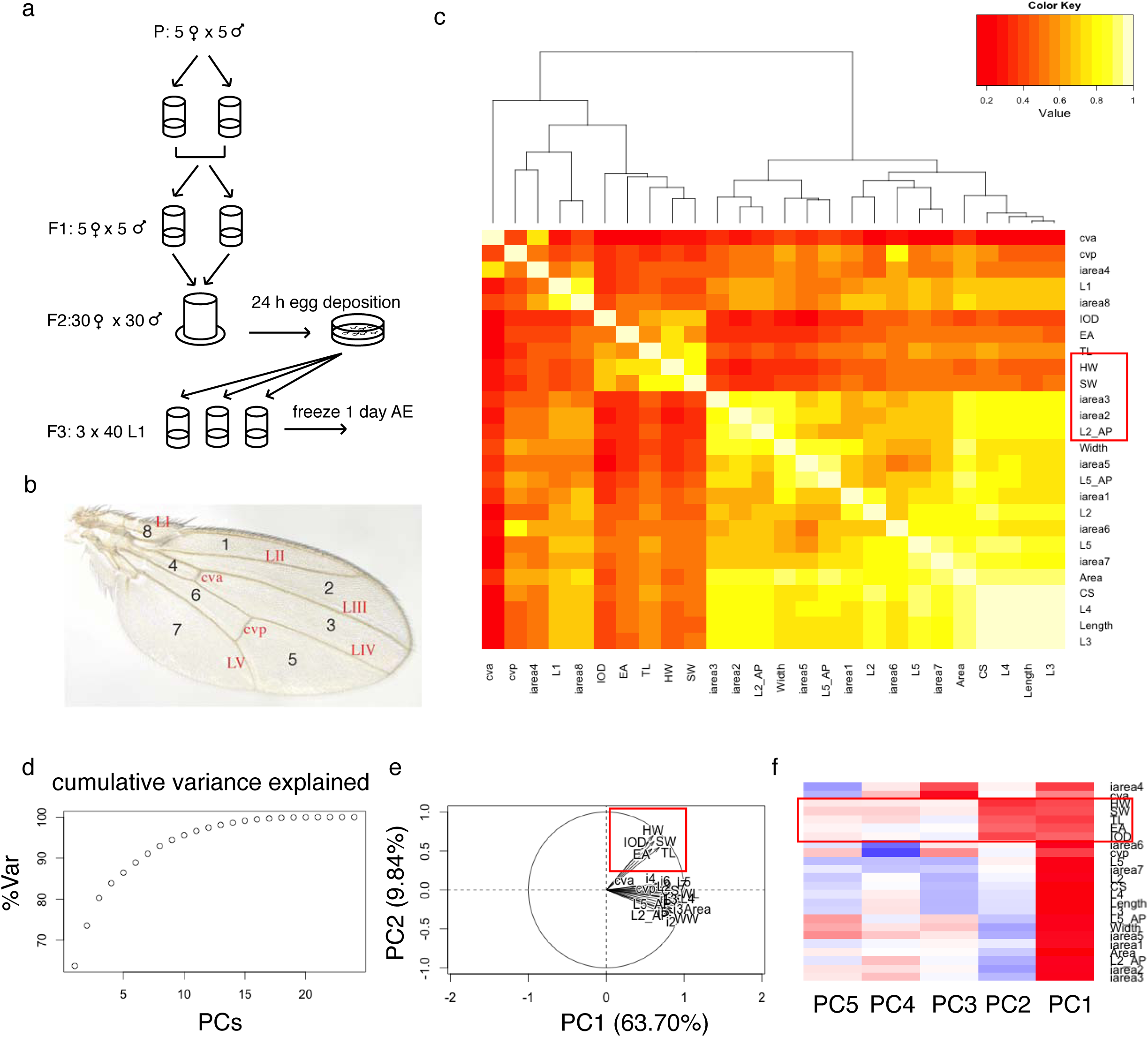
Analysis of 26 size traits in the DGRP. (a) Standardized *Drosophila* culture conditions for the quantification of morphometric traits. The protocol extends over three generations and efficiently controls known covariates of size, such as temperature, humidity, day-night-cycle and crowding. Additionally, effects of other environmental covariates, such as intra-vial environment, light intensity and incubator position, are randomized. (b) Illustration of the wing features. L2_AP and L5_AP are not illustrated; they comprise the area between the AP boundary and L2 or L5, respectively, and serve as measures for the size of the anterior and posterior part of the wing. (c) Genetic correlation between morphometric traits in females. Two modules of higher correlation are clearly visible (bright yellow): one of almost all wing features and one comprising all head/thorax traits. (d) Cumulative variance explained in female data by increasing number of principal components. (e) Variables factor map. PC1 and PC2 split the data into two groups. (f) Correlation between PCs and traits. PC1 reflects a general size component and PC2 is highly correlated with head/thorax traits, effectively splitting the data in two groups.

Given the observed redundancy of the phenotypes, we chose only one trait from each high-correlation module for further in-depth analysis: centroid size (CS, reflecting growth processes in the wing disc), and interocular distance (IOD, reflecting eye disc growth), respectively (Fig. 2a,b). IOD showed the lowest genetic correlation with CS of all head/thorax traits (0.46 in females and 0.51 in males). Interestingly, the allometric coefficient b describing the relationship *CS = a*X^b^* (where *X* = IOD or TL) varied substantially between lines, from near independence (b=0) to hyperallometry (positive allometry b>1) (S2 Fig., S1 Table). We observed extensive phenotypic and genetic variation in both phenotypes (Fig. 2a, b, S1 and S2 Tables), which was reflected in the substantial broad-sense heritabilities 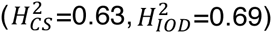. Furthermore, both traits showed significant genetic variation in sex dimorphism but similar heritability estimates for males and females and high cross-sex genetic (*r*_*MF*_) and phenotypic correlation (S1 Table). 15% of phenotypic variance in centroid size could be attributed to raising flies on different food batches, which only differed by the day on which they were prepared (according to the same protocol) (S1 Table). Though nutrition is a well-studied size-determining factor (52), we were surprised that even such small nutritional variation could have substantial phenotypic effects. Although the environmental effect of food batch was markedly lower for IOD (3%), we used batch-mean corrected phenotypes in all subsequent analyses to remove this effect.

**Figure 2.**
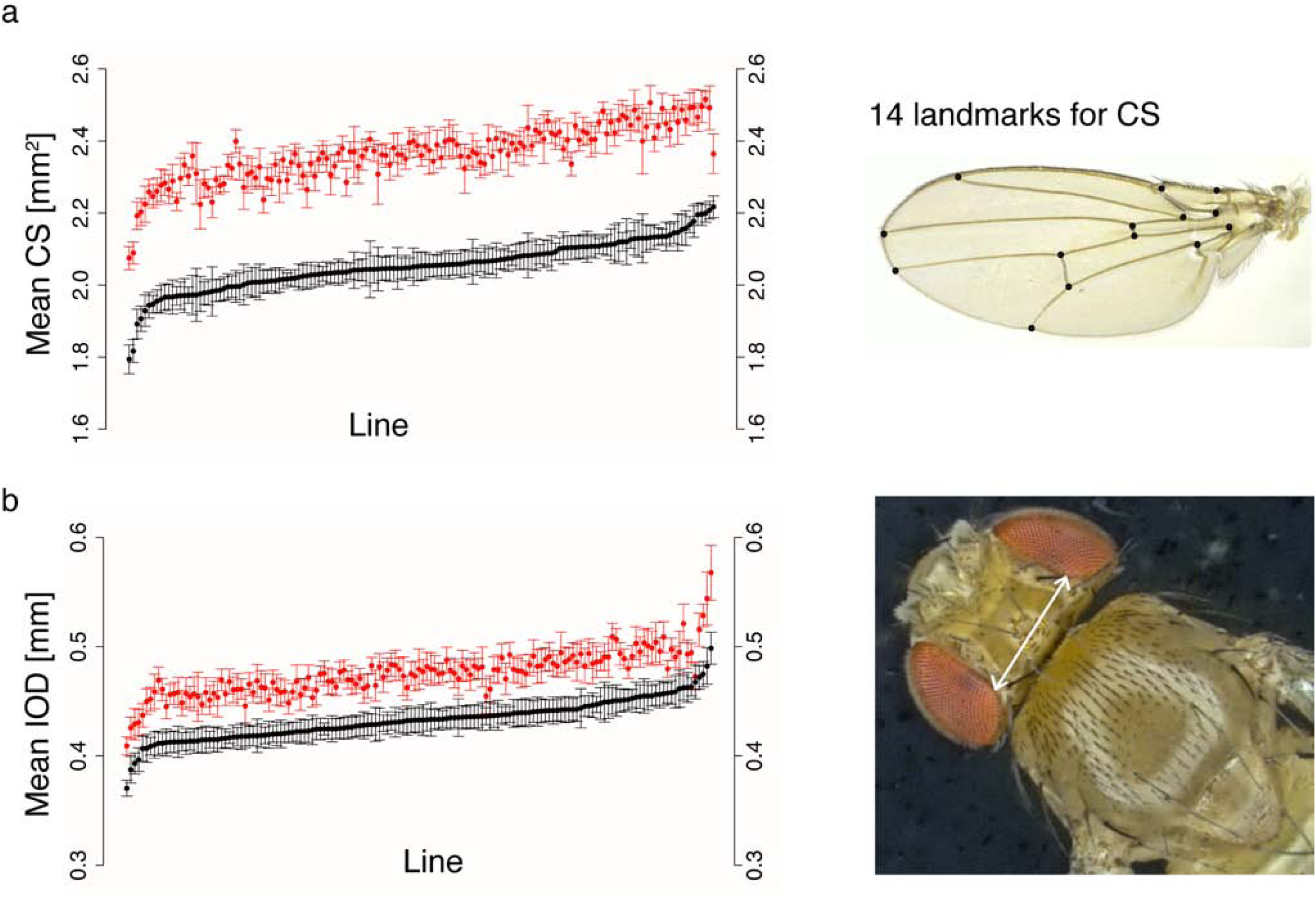
Phenotypic variation in the DGRP for two size traits. Plots show mean phenotypic values for (a) centroid size and (b) interocular distance. Each dot represents the mean phenotype per line of males (black) and corresponding females (red), with error bars denoting one standard deviation. Lines are ordered on the *x*-axis according to male trait value, from lowest to highest, meaning that the order of lines is different for each plot. Raw phenotypes and line means are listed in S2 Table. To the right are illustrations of both measures.

### Single-marker and gene based GWAS identify novel loci associated with size variation

To identify common loci contributing to size variation in *Drosophila*, we performed single marker GWA analyses (53) for 1,319,937 SNPs for a wing disc derived (CS) and an eye disc derived (IOD) size measure using Fast-LMM. This association method uses a linear mixed model to capture confounders such as population structure and cryptic relatedness. As the genetic correlation between CS and IOD was moderate (0.46 and 0.51 for females and males, respectively), we expected to map both shared and trait-specific SNPs. To find loci that specifically affect variation in wing size unrelated to the overall organismal size variation we constructed an additional phenotype (rCS) that had the effect of IOD on CS removed via regression. In addition to the effect of the food batch, two cosmopolitan inversions, *In(2L)t* and *In(3R)Mo*, were correlated with both CS and IOD and we addressed their effect on size by modeling their presence in the homozygous state (S2 Fig.), yielding the inversion-corrected phenotypes CS_IC_ and IOD_IC_. *In(3R)P*, which is known to be correlated with *Drosophila* size (19), was only homozygous in one line; therefore, we could not estimate its effect on size.

Only for one trait (IOD in females) did we observed significantly associated SNPs when applying a stringent Bonferroni corrected *p*-value threshold of 3.8×10^-08^. However, their significance dropped below the genome-wide level (*p*_*min*_=3.94×10^-06^) when we applied GWAS on rank-normalized IOD, which is probably due to an outlier line (>4SD) in the minor allele class of these SNPs. Overall, the *p*-values between normalized and non-normalized GWAS showed good correlation (S3 Fig.).The six SNPs were all located in a cluster on chromosome *2L (2L: 12’805’398 – 12’806’812)*, 12-13kb upstream of the gene encoding the EGFR pathway regulator *kek1* (Fig. 3a,c). Three more SNPs in this locus were annotated to *kek1*, but did not survive Bonferroni correction. All nine SNPs formed a haplotype, with lines having either all minor or all major alleles of these SNPs, and the minor allele haplotype was associated with an increased IOD (Figs. 3b,d, S3 Fig.). 198 SNPs are located in the region approximately 20kb upstream of *kek1*. This region showed high conservation between species (DGRP Freeze 2 genome browser, http://genome.ucsc.edu) and several blocks of higher LD are formed across it (Fig. 3b, S3 Fig.), which could be attributable to its proximity to *In(2L)t* (*2L: 2’400’000 – 12’900’000*). However, none of the lines with the minor allele haplotype was either homo-or heterozygous for this inversion, and they were distributed across all four food batches (S3 Fig.). Interestingly, a noncoding RNA, *CR43818*, is in the 20kb upstream region of *kek1*, and the region was spanned by the intron of *CG9932*, a poorly characterized gene that interacts genetically with *Bx* and *Chi* during wing development (54). Clearly there are signs for functionality of this locus, and several good candidates for causal variants. Further experiments are required to elucidate the molecular mechanism of this association and whether this may even be connected to *In(2L)t*.

As QQ-plots showed a departure from uniformity for *p*-values below 10^-05^ (S4 and S5 Figs.), and this threshold has been used for reporting associations in previous DGRP studies (48) we picked candidate loci using this nominal significance threshold for hypothesis generation and functional validation. The corresponding *q*-values for each SNP are listed in S3 Table. This yielded between 31 and 51 SNPs for females and between 17 and 36 SNPs for males, with little overlap between top associations and moderate correlation of overall SNP ranks between sexes (S3 and S4 Tables; S6, S7 and S8 Figs.), consistent with significant sex by line variances and departure of the cross-sex genetic correlations from unity in the quantitative genetic analyses.

**Figure 3.**
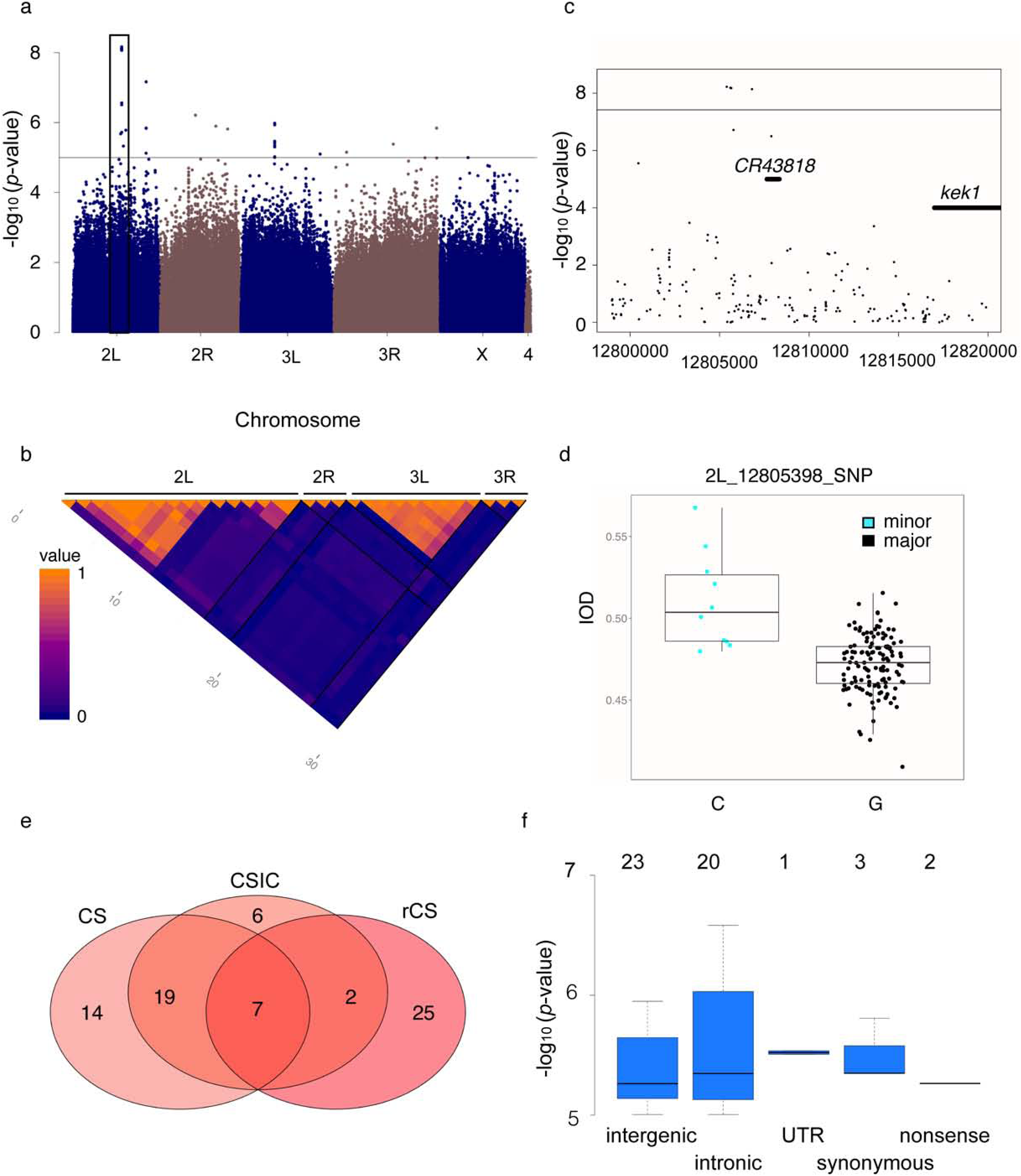
Genome-wide association of size traits. (a) Manhattan plot of the SNP *p*-values from the IOD GWAS in females shows that nominally associated SNPs are distributed over all chromosomes. Negative log10 *p*-values are plotted against genomic position, the black horizontal line denotes the nominal significance threshold of 10^-05^ and the black box marks the location of the cluster of Bonferroni-significant SNPs upstream of *kek1 on 2L*. (b) Correlation between SNPs nominally associated with female IOD. The cluster of Bonferroni-significant SNPs on *2L* shows high correlation among individual SNPs over a larger region, whereas most other SNPs except a few in a narrow region on *3L* represent individual associations. Blue = No correlation, orange = complete correlation. Pixels represent individual SNPs and black lines divide chromosomes. (c) Locus zoom plot of the region 20kb upstream of *kek1* hat harbors the genome-wide significant associations. The black horizontal line denotes the genome-wide significance threshold (*p*=3.8×10^-08^) and the locations of *kek1* and the ncRNA *CR43818* are marked by broad black lines. (d) Lines with the minor allele genotype at the most significantly associated locus have a larger IOD than lines with the major allele. (e) Overlap in the number of nominally associated SNPs for different wing traits in females. The overlap is bigger between the absolute wing size phenotypes and only a few SNPs are candidates for all traits. (f) Nominally associated SNPs are most abundant in the intergenic space and in regulatory regions. Boxes show the distribution of negative log10 *p*-values of the SNPs nominally associated to rCS in females among site classes. Numbers of SNPs belonging to each site class are denoted above the boxes. As a SNP can fall into multiple classes, the sum of SNPs from all site classes is higher than the total number of nominally associated SNPs.

Correcting for the segregation of polymorphic inversions generally enhanced the power of the GWA analyses, as is evident by more loci reaching nominal significance. Nevertheless, the majority (65-86%) of SNPs from the GWA analysis with uncorrected trait values remained candidates in the GWA analysis with corrected phenotypes. Somewhat surprisingly, despite the significant genetic correlation between CS and IOD, no candidate SNPs were shared between these phenotypes (Fig. 3e, S4 Table). In both sexes, approximately one-third of top SNPs was shared between the absolute and relative CS GWA analyses, suggesting variation in relative versus absolute organ size may be achieved through genetic variation at both shared and private loci.

Nominally associated variants predominantly mapped to intergenic regions, but were nevertheless enriched in gene regions (*p*<0.001, hypergeometric test) (Fig. 3f, S5 Table), demonstrating that associations were not randomly distributed across the genome. For gene-level analyses we determined candidates for each phenotype as genes having a nominally significant SNP in or within 1kb of their transcribed region, yielding a total of 107 genes over all phenotypes. Only the candidate gene sets for rCS were enriched for STRING curated interactions and only the candidate list for CSF was enriched for functional categories (positive regulation of Rho signal transduction and melanotic encapsulation of foreign target), though growth was among the top categories for CSM_IC_ (FDR corrected *p*=0.08) (55). Given the large number of genes already known to play a role in growth control we were surprised that only few canonical growth genes contained or were close to nominally associated SNPs. Exceptions included several SNPs near or in the genes coding for Ilp8, TOR and EGFR pathway components and regulators of tissue polarity and patterning. However, some SNPs that narrowly missed the candidate reporting threshold localized to further growth regulatory genes, such as the Hippo pathway components *ex* and *wts*.

The small number of canonical growth pathway genes detected might be explained by the lack of SNPs with large effects in these genes, which is plausible considering the essential role of many growth regulators. We therefore wanted to test whether the combined signal of SNPs with small effects (each too small to reach significance on its own) across known growth genes might be significant. To this end we determined gene-based statistics using the sum of chi-squares VEGAS method (56), which computes a *p*-value for each gene considering all SNPs within a gene while correcting for gene length and linkage disequilibrium between SNPs. None of the genes reached genome-wide significance (*p*<3.75×10^-06^) (S6 Table). The overlap between the 20 top scoring genes from this analysis with our GWA candidate genes was small for each individual phenotype and even when combining the VEGAS analyses from all phenotypes only 11 of our 97 VEGAS top scoring genes contained a SNP that reached significance on its own in one of our GWA analyses. We did not find GO or interaction enrichment (55) and as in the individual GWA analyses, top candidates were largely novel with respect to growth control.

### Functional validation of candidate genes reveals novel regulators of size

We selected a subset (41% to 69%) of candidates identified by each of our six wing size GWAS (CS, CS_IC_ and rCS in both sexes) for functional validation by tissue-specific RNAi. A total of 64% to 79% of tested genes had significant effects on wing area (*p*<0.001, Wilcoxon rank sum test, S7 Table, Fig. 4a, S9 Fig.). We achieved similar validation rates for gene-based candidates. In contrast, only 42% of a set of 24 randomly selected genes had significant effects on wing size in females (S7 Table). The overall proportion of validated candidates versus random genes was significantly different (*p*=0.02, Fisher’s exact test) and Wilcoxon test *p*-values showed different distributions between candidate and random knockdowns (*p*=0.02, Wilcoxon test, S10 Fig.). This combined evidence suggests an advantage in power for identifying growth regulators by GWA over randomly testing genes. The validated candidates constitute 33 functionally diverse novel growth regulators (S11 Fig.).

**Figure 4.**
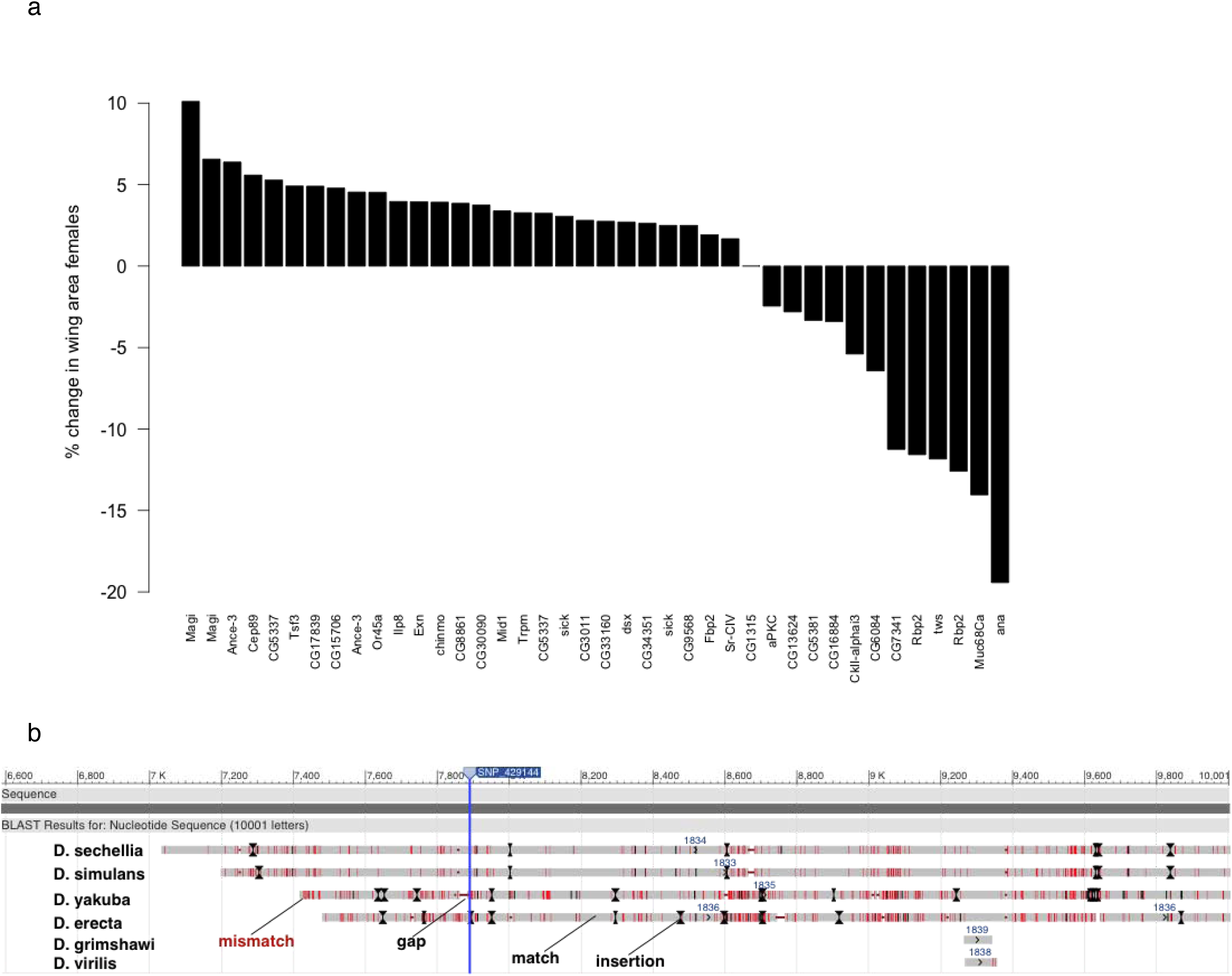
Associated SNPs overlap 33 functionally diverse novel candidate genes for wing size determination and localize within putative enhancer elements. (a) Validated genes in females. Bars show the percent change in median wing area compared to *CG1315* RNAi upon wing-specific knockdown of candidate genes. Only the lines yielding a significant wing size change (*p*<0.001, Wilcoxon rank sum test) are depicted. (b) Alignment of the 2kb region on chromosome arm *2L* upstream of the *D. melanogaster ex* locus that shows sequence conservation across *Drosophila* species. The position of the SNP is indicated by the vertical blue line. The *D. melanogaster* sequence is represented by the dark grey bar at the top (“Sequence”). The respective sequences of each compared species are represented below. Light grey regions are matches to the *D. melanogaster* sequence, red regions are mismatches, gaps in the alignment are denoted by horizontal red lines and insertions by black lines and arrows.

### Two-locus association reveals novel interactions

To place novel genes within the network of known growth pathways, we next performed tests for two-locus associations (57) to CS_IC_, IOD_IC_ and rCS in both sexes with SNPs in 306 growth genes as focal SNPs (S8 Table). This gene list was combined from genes listed as influencing wing development (The Interactive Fly, http://www.sdbonline.org/sites/fly/aimorph/wing.htm), commonly known growth genes from the IIS/TOR, EGFR and Hippo pathways, and growth regulators identified in screens by our group. This list is not comprehensive but should serve as a rough framework for the most relevant growth pathways. Overall, 15 interactions reached Bonferroni-corrected significance (*p*<7.9×10^-13^), but we observed none of our GWA candidates among the significant epistasis partners. Generally, more interactions reached genome wide significance in males than in females. The most significant interaction (CSM_IC_, *p*=5.79×10^-15^) occurred between *mask*, a positive regulator of JAK/STAT signaling (58) and *tutl*, a JAK/STAT target gene during optic lobe development (59) (Fig. 5). Furthermore, among the top five interactions we found one between *nkd,* a downstream target of *Dpp* (60), and the tyrosine phosphatase *Ptp99A* (CSF_IC_, *p*=8.79×10^-14^), which has been shown to interact with InR and the Ras signaling pathway (61,62). Furthermore, though we detected none of the significant interactions on DroID (63,64) *mask* and *tutl*, and *PtP99A* and *nkd* shared more DroID interactors than 95% of all possible pairs of the 32 genes involved in the significant interactions. Due to their already known growth-related functions we consider the interactions between these genes prime candidates for future functional validation.

**Figure 5.**
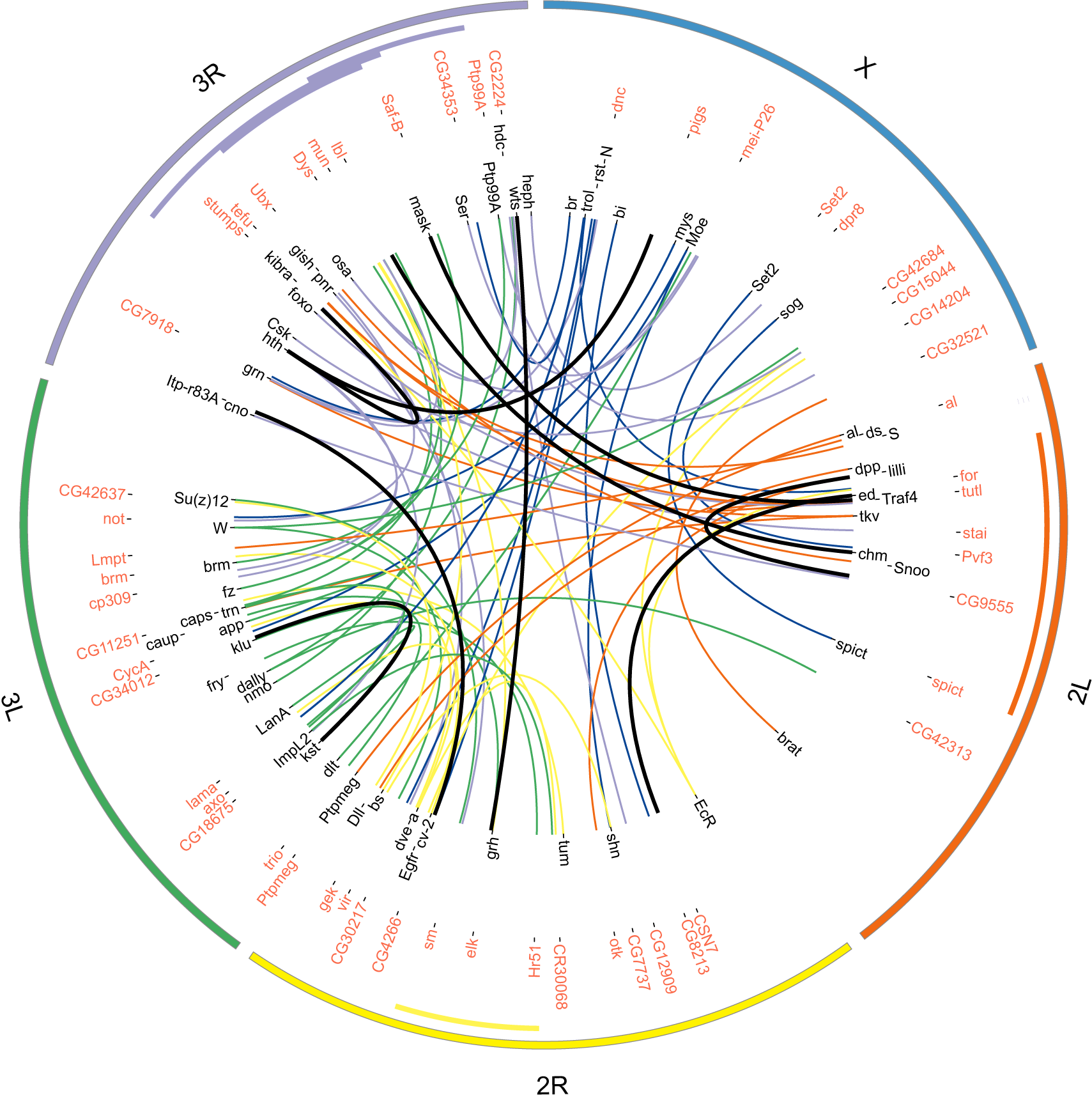
Pairwise interactions between focal genes and DGRP SNPs for male wing size (rCSM). The plot shows the focal genes annotated in black and the interactors in red. Interaction lines are colored according to the chromosome the focal gene is located on and the thick black lines denote Bonferroni-significant interactions. The outer circle demarks the chromosome arms (*2L* = orange, *2R* = yellow, *3L* = green, *3R* = purple, *X* = blue). The colored bars inside the inner circle demark the locations of cosmopolitan inversions (orange: *In(2L)t*; yellow: *In(2R)NS*; purple: *In(3R)K*, *In(3R)P*, *In(3R)Mo*).

To investigate whether our GWA candidates or genes from the ‘previously known’ catalog would be enriched further down the list, we lowered the stringency for reporting interactions to a discovery threshold of *p*<10^-09^. Counting only those interactions where the interacting SNP lay in or within 1kb of a gene (S8 Table, total 1,353 interactors across all phenotypes) we found enrichment for development, morphogenesis and signaling categories (Bonferroni corrected *p*<0.001) (55), which supports a role of these genes in growth control. Notably, the rCSM list (Fig. 5) was additionally enriched for genes involved in regulation of metabolic processes. Among them were 73 of the 306 genes in the ‘previously known’ catalog and 35 of our 107 overall GWAS candidates. However, these overlaps did not reach significance (*p*-value of 0.46 and 0.22, respectively). We next asked whether the candidate gene sets identified by normal GWAS and the epistasis approach were nevertheless biologically related to each other. To this end we used the STRING database (55), which revealed that the number of observed curated interactions between the two gene sets was much larger than expected by chance (*p*<<0.001). Analyzing pairwise interactions may thus help to place genes into pre-established networks.

### Intergenic SNPs are preferentially located in regions with enhancer signatures and overlap lincRNA loci

Intergenic SNPs may be functional by changing the sequence of more distant regulatory elements or noncoding RNAs. We therefore tested whether intergenic GWAS candidate SNPs located to putative functional regions. We found enrichment (*p*<0.01, hypergeometric test) of SNPs lying in regions with H3K4Me1 or H3K27Ac, epigenetic signatures of active enhancers (S5 Table) (65), and in lincRNA loci (66), which have been implied in developmental regulation and are often enriched for trait-associated loci (67). Though only loci associated with IOD in females were enriched for SNPs localizing to lincRNA loci, we found one SNP lying in a lincRNA among the top variants for rCSF and IODF_IC_ (S5 Table).

A SNP 2kb upstream (position *2L*: 429144) of the Hippo pathway regulator *ex* narrowly missed the reporting threshold (*p*=1.7×10^-05^, CSM). However, its genomic location suggests this variant could affect a novel regulatory region for this gene. The region surrounding it was annotated with the enhancer methylation signatures H3K4Me1 and H3K27Ac and had assigned state 4 of the 9 state chromatin model suggestive of a strong enhancer (65,68). Further annotations included H3K9Ac, a mark of transcriptional start sites, histone deacetylase binding sites and an origin of replication. To further assess functionality, we investigated whether the sequence around this SNP was conserved across taxa by performing multiple sequence alignment using BLAST (69) (S9 Table). Indeed, the region immediately upstream of the *D. melanogaster ex* gene showed high similarity to ∼3kb regions slightly more upstream of *expanded* orthologs in the genomes of *D. sechellia, D. yakuba* and *erecta* (S9 Table, Fig. 4b). This combined evidence suggests a functional region immediately upstream of the *D. melanogaster ex* gene, but additional experiments are required to corroborate functionality and to establish an involvement in growth control and a mechanism for influencing size.

### Human orthologs of candidate genes are associated with height, obesity and a variety of other traits

To investigate conservation to humans and further elucidate putative functions of candidate genes, we searched for orthologous proteins in humans. We found human orthologs (70) for 62 of our 107 GWA candidate genes, seven of which had a good confidence ortholog (score ≥ 3) associated with height, pubertal anthropometrics or growth defects (S10 Table). Furthermore, the GWAS candidates were enriched for human orthologs associated with height in a recent meta-analysis (*p*= 0.001) (10). The evidence for an involvement in growth control from GWAS in both organisms and experimental support from validation in *Drosophila* corroborates a biological function of these genes in the determination of body size.

## DISCUSSION

We applied several GWA methods to developmental traits that have been extensively studied by single gene analyses in *Drosophila* as a complementary approach for identifying loci underlying size variation. Our single-marker GWAS revealed only one SNP cluster close to the known growth gene *kek1* to be significantly associated with body size when using a conservative Bonferroni correction. Yet, in contrast to human GWA analyses, which require independent replication, we exploited the fact that our model organism is amenable to direct validation strategies and tested candidates corresponding to a much lower significance threshold of 10^-05^ for an involvement in size determination. Using tissue-specific RNAi, we validated 33 novel genes affecting *Drosophila* wing size. Nominally significant intergenic associations were preferentially located in regions with an enhancer signature and overlapped lincRNA loci. A SNP upstream of the *expanded* locus was in an evolutionarily conserved region, indicating the presence of a putatively functional element. A two-locus epistasis screen identified several genome-wide significant interactions between known growth genes and novel loci, showing that targeted epistasis analysis can be used to extend existing networks. Our study shows that despite limited statistical power, insights into the genetic basis of trait variation can be gained from analyzing nominal associations through functional and enrichment analyses and performing targeted locus-locus interaction studies.

### Single-marker and two-locus GWA of size

Our study adds 33 novel growth genes and 15 genomic loci that may interact with known growth genes to the extensive number of loci already implied in size regulation from single gene studies. That only a few *bona fide* growth genes were among the nominally significant candidates could be due to selection against functional variation in natural populations and/or during subsequent inbreeding and/or that the effect sizes of SNPs in these genes are too small to be detected in the DGRP. Though it has been shown for other phenotypes that the identified loci seldom overlap between mutational and GWA approaches, we expected a higher overlap for size as this trait has been exceptionally intensively studied in *Drosophila* and as a result we have extensive prior knowledge on the underlying genes. Our validation of a substantial number of novel genes underscores the complementarity of the GWA approach to classical genetics and highlights the importance of probing natural variants. However, future studies would benefit from utilizing bigger population sizes in order to improve statistical power and from investigating populations with different geographic origins, to address population specificity of associated variants.

With the exception of the *kek1* cluster, all the most significant wing size associations mapped to putative novel growth genes: *CG6091*, a de-ubiquitinating enzyme whose human ortholog has a role in innate immunity; *CG34370*, which was recently identified in a GWA analysis of lifespan and lifetime fecundity in *Drosophila* (71); and, surprisingly, *dsx*, a gene well characterized for its involvement in sex determination, fecundity and courtship behavior. *dsx* showed a significant effect on wing size in both sexes, *CG6091* was only validated in males despite reaching a smaller *p-*value in the female GWAS, and *CG34370* did not show a significant wing size change in either sex. Due to the obvious limitations of RNAi as a validation approach for SNPs we think it important to investigate the roles of *CG6091* and *CG34370* in growth control by other approaches before discarding them as false associations. As genes affecting growth also impact on general and reproductive fitness of organisms it is not surprising that most of the candidate variants in or close to genes lie in regulatory regions, potentially modulating splicing, RNA turnover or RNA/protein abundance. Our data support the general notion that intergenic SNPs can impact phenotypes, either by affecting transcript abundance of protein coding genes (e.g. through distal enhancer elements) or via noncoding RNAs, which have been shown to regulate many biological functions including cellular processes underlying growth (72,73,74).

There are few nonsense and missense SNPs among our candidates (one and six, respectively); these variants are prime contenders for effects on protein function. However, confirming such effects requires testing the SNP in an isogenic background. Knockdown of most candidate genes resulted in a small change in median wing size (-19.4% to 10.1%), indicating a redundant or mildly growth enhancing or suppressing role in this tissue, which may explain why they were not discovered by classical mutagenesis screens. However, larger effects might be observed upon ubiquitous knockout, knockdown or overexpression.

Epistasis analysis revealed 15 loci showing Bonferroni-significant interactions with SNPs in previously known growth genes, demonstrating the usefulness of this approach for extending existing biological networks. In addition to the interactions described in the main text, we note there is an interesting interaction between *eIF2D* (*ligatin*) and *SNF4agamma* (75). Furthermore, we found putative biological interactors for several GWA candidates among the top interactions that did not reach genome-wide significance e.g. *Lar* with *InR*. Lar can phosphorylate InR (61), so polymorphisms at these two loci could act synergistically to modulate InR activity. The enrichment of annotated interactions between our GWA candidates and epistasis partners shows that different analyses yielding different top associations uncover common underlying genetic networks. A similar combinatorial approach has been successful in study using the DGRP (33), underscoring that combinatorial approaches can help placing candidates from different analyses into a joint biological network, and provide a basis for further hypothesis driven investigation of the roles and connectivity of novel and known genes.

### The *kek1* locus

The region upstream of *kek1* is the only locus where a larger genome region shows association. The minor allele frequency of this haplotype was between 4.7% and 6.2% (present in 7-10 lines). We explain the large effect of this locus by the fact that most lines with the minor alleles of the significant SNPs have an IOD that exceeds the 75^th^ percentile of the IOD distribution in lines with the major allele. Obviously the effect may be overestimated due to relatively few lines having the minor allele, and due to effects of the genetic background. Nevertheless, the region contains some prime contenders for an effect on size: the SNPs lie in a region that could serve as a regulatory element for *kek1* or the poorly characterized *CG9932* that has been implied in wing disc development in another study (54). Furthermore, an uncharacterized noncoding RNA lies close by and could be linked to the SNP haplotype, due to generally higher LD in this region. Also, the region is spanned by *In(2L)t*, which itself shows association to size, and the observed effect could be due to this inversion. None of the lines with the minor allele haplotype had any copy of the inversion, which agrees with the observation that *In(2L)t* homozygous flies are relatively smaller than lines without the arrangement. On the other hand, only two lines in our dataset were homozygous for *In(2L)t* (DGRP_350 and DGRP_358), leaving many of the lines with the major allele with none or only one copy (nine lines) of the inversion, and we corrected for its presence, so we would not expect this to cause the association.

### Biological roles of novel growth genes

Apart from expected processes like signaling, transcription, translation and morphogenesis, we validated genes involved in transmembrane transport, planar cell polarity (PCP), metabolism and immunity. A total of 24 single marker or two way interaction candidate genes from our GWA analyses were discovered to be enhancers or suppressors of major growth pathways in another study (76, S11 Table) and 15 were associated with nutritional variation in *Drosophila* (77), supporting their role in growth control. Importantly, the enrichment of GWA candidates for genes associated with human height not only supports their role in growth but also shows that results of GWA studies in *Drosophila* can be translated to orthologous traits and genes in humans. We do not see a large overlap between our candidates and the candidates identified by artificial selection on body size by Turner *et al.* (16), who identified many classical growth genes. We explain this discrepancy by the different methods to recover underlying genetic loci. Selection can enrich for rare alleles, while these cannot be probed in GWA. If these also had large effects, they would be strongly differentially selected for in the Turner study. In contrast, we would expect large effect alleles of canonical growth genes to be mostly rare in the DGRP lines, and thus not evaluated in the GWA. Furthermore, Turner *et al.* used a population from California, which likely has a different allele composition from our North Carolina population. A minor factor may be the rather coarse and general size measure used by Turner *et al.* Sieving selected for generally bigger flies, with no distinction between flies with bigger wings, heads, thoraxes, or legs that obstructed passage through the sieve. As growth of individual body parts is controlled by both systemic and organ intrinsic factors (1-4), and looking at overall size likely identifies the systemic, general factors, this could explain some of the discrepancy. We do not identify either *InR* or *smp-30* (23,24). Both alleles have a relatively large effect. The *smp-30* allele was identified in a population of different geographic origin and could thus not be present in the DGRP lines. The *InR* allele is an indel, and we only analyzed single nucleotide polymorphisms in this study. Although both genes likely contain further polymorphisms that are present in the DGRP, their effects may be far smaller and thus not detectable given the restricted size of the DGRP. In terms of chromosomal loci, we do identify SNPs on *3R*, several of them located in the region spanned by *In(3R)P (3R: 12*,*257*,931 – 20,569,732). Also, we see a suggestion of *In(3R)P* correlating with size, but the inversion was present in too few lines of our dataset to reliably estimate its effect in a model.

Most of the loci we validated have not been previously linked with growth in *Drosophila*. The yeast ortholog of *Mid1* (validated only in females but stronger association in males), is a stretch activated Ca^2+^ channel with a role in the polarized growth of mating projections (78,79). As mechanical tension plays a role during growth of imaginal discs, this channel could act in translating such signals to intracellular signaling pathways via the second messenger Ca^2+^. The human ortholog of another candidate, the transmembrane channel *Trpm* (associated and validated only in females), showed association with anthropometric traits during puberty, indicating a role during the postnatal growth phase. The mucin *Muc68Ca*, identified in the top 20 of the gene-based association to rCSF, showed one of the largest knockdown effects (14 and 15% reduction in size in females and males, respectively). Mucins form a protective layer around vital organs, and the expression pattern of *Muc68Ca* in the larval midgut concurs with a putative effect on growth via the control of intestinal integrity (80).

A dual role in PCP, the establishment of cell polarity within a plane in an epithelium, and growth control has been shown for many genes, which regulate these two processes via distinct but coordinated downstream cascades (81,82). *Lar* (no significant effect on size in validation), *aPKC,* the Fz target Kermit (83) and the motor proteins Dhc64C (not included in validation but contained a nonsense SNP reaching nominal significance) and Khc-73 (validated in males but stronger association in females, though with positive effect size), whose human ortholog is significantly associated with height, are implied in PCP establishment. Kermit and motor proteins act downstream in the PCP cascade and likely have specialized roles for this process, but PCP can itself impact on growth, as proper establishment of polarity provides the orientation of cell division, and loss of a PCP component in zebrafish causes a reduction in body length (84). Interestingly, *kermit* was a candidate interactor of EGFR, which acts in a combinatorial manner with Fz signaling in PCP (85), providing a biological basis for this interaction.

Metabolic genes are prime candidates for improving our understanding of growth, which depends on the amount of energy and precursors available for biosynthesis, and thus to metabolic coordination. The recent findings that the growth and PCP regulator Fat can couple growth and metabolism and mitochondrial proteins can causally affect growth pathway activity (86) underscore the importance of metabolic coordination. A missense SNP in the validated candidate *Cep89*, a gene involved in mitochondrial metabolism and growth in *Drosophila* and humans (87) was associated with most wing phenotypes. Elucidating the function of *Cep89* and other validated candidates with putative roles in metabolism, e.g. *CG3011*, *CG6084* and *Fbp2,* whose human ortholog has been linked to growth defects and cancer (*e.g.* 88), may provide further insight into this coordination.

### Sexual dimorphism of size

Of the top 100 SNPs for each trait only 25% - 43% are shared between the sexes, a surprisingly small overlap given the high genetic and phenotypic correlations between sexes. In some cases the SNPs still lie in the same gene, implying that this gene differentially affects size in both sexes, but the responsible SNP is different. In other cases, we only detect associated SNPs for a gene in one sex. Here, the gene may affect size in both sexes but genetic variation in this gene affects size differentially in only one sex. As we have a low powered study this is only a hint and these results need to be further analyzed in a bigger population or by allele replacements using e.g. CRISPR/Cas9 to be corroborated. Unfortunately we cannot conclude anything about sex-specificity of our variants from the knockdown results. A knockdown is a very different perturbation from the effects of, for example, a regulatory variant. The knockdowns are performed in a different background than the one the association was discovered in, and they have much larger effect on the levels of a gene than a regulatory variant. So even though RNAi on a gene might show effects in males and females it does not exclude that different alleles of a SNP in this gene only affect wing size differentially in one sex.

Interestingly, an intronic SNP in the sex determination gene *dsx* had the lowest *p*-value in the female relative wing size GWAS but had a smaller effect size in males. Dsx is a transcription factor with sex-specific isoforms, and has many targets with sex- and tissue specific effects (89). We also observed sex-specificity for the genome-wide significant two-locus interactions. Considering that males use their wings to produce a courtship song that is instrumental for mating success, it may well be possible that selection pressure is different for male and female wing size or wing size in relation to other body parts. Indeed, the selection response for wing length seems to be more constrained in males (90). In our dataset, wing length is highly correlated with CS, our main wing size measure. Menezes *et al.* observed males with more elongated wings but also smaller males had the highest mating successes (91). These studies and our data suggest there may be subtle differences in the genetic networks underlying size determination in males and females in natural populations, a possibility that is neglected in single gene studies and thus would be worthwhile exploring.

### Conclusions and future perspectives

Growth control has been well studied, particularly in *Drosophila,* where many genes and pathways affecting growth have been documented by mutational analyses. However, such screens are far from saturation and do not scale well to investigating effects of combinations of mutations. Here we took advantage of naturally occurring, multifactorial perturbations genome-wide to identify novel genes affecting growth and to place them in genetic interaction networks. Rather than deepening our understanding of growth control, the identification of ever more growth regulators raises new questions about how all these loci interact to govern growth. The challenge for the future will be to shift our focus from studying genes in isolation towards investigating them in the context of developmental networks, and to assess the effects of network perturbations on intermediate molecular phenotypes of transcript, protein and metabolite levels.

## MATERIALS AND METHODS

### *Drosophila* medium and strains

Fly food was prepared according to the following recipe: 100 g fresh yeast, 55 g cornmeal, 10 g wheat flour, 75 g sugar, 8 g bacto-agar and 1 liter tap water. Experiments were performed with 149 of the DGRP lines. RNAi lines used are listed in S7 Table.

### Standardized culture conditions

Lines were set up in duplicate vials, with five males and five females per vial. After seven days, the parental flies were removed. From the F_1_, five males and five females were put together in duplicate vials and discarded after seven days of egg laying. From the F_2_, thirty males and thirty females were mated in a laying cage with an apple juice agar plate plus a yeast drop as food source and allowed to acclimatize for 24 hours. A fresh plate of apple juice agar plus yeast drop was then applied and flies were left to lay eggs for another 24 hours. From this plate, F_3_ L1 larvae were picked with forceps and distributed into three replicate vials, with 40 larvae per vial. The food surface in the vials was scratched and 100*μ*l of ddH_2_O added prior to larvae transfer. The adult F_3_ flies were pooled from the three vials and frozen at -20°C approximately 1-2 days after eclosion. The whole experiment was performed in a dedicated incubator (DR-36VL, CLF Plant Climatics GmbH) with a 12-hour day-night cycle, constant humidity of 65 - 68% and constant temperature of 25.5°C +/- 1°C. Vials were shuffled every two days during the first and second round of mating but left at a fixed position in the incubator for the duration of the development of the F_3_ generation.

For the parental generation, lines were all set up on the same day on the same food batch. For the F_1_ matings, different food batches had to be used due to different developmental timing of the lines. F_2_ matings were set up using the same batch of apple agar plates and yeast for all lines. F_3_ larvae were distributed on four different food batches and the batch number was recorded for each line.

The control experiment (S1 Table) was performed using the same procedure as above, except that the same food batch was used for all flies of a generation. We used the DGRP lines DGRP_303, DGRP_732, DGRP_721 and DGRP_908 for this experiment because they had comparable generation times and set up ten replicates of each of these lines according to the standardized culture conditions.

### Phenotyping and morphometric measurements

Depending on the number of flies available, between five and twenty-five flies per sex and line were measured for the dataset (median 25 flies per sex and line, mean 23 (CS_females_, CS_males_, IOD_males_) and 24 for IOD_females_; exact numbers are given in S2 Table). For the experimental generation we distributed a total of 19,200 larvae in four batches spaced throughout 1.5 weeks according to developmental timing, and the final dataset consisted of morphometric data of 6,978 flies, 3,500 females and 3,478 males. For the control experiment we phenotyped 25 flies per replicate, sex and line, resulting in a total of 2,000 flies (1,000 males, 1,000 females). Flies were positioned on a black apple agar plate and photographed using a VHX-1000 digital light microscope (KEYENCE). Morphometric body traits were measured manually using the VHX-1000 dedicated measurement software. If intact the right and otherwise the left wing was removed and mounted in water on a glass slide for wing image acquisition. Morphometric measurements were extracted from the wing images using WINGMACHINE (92) and MATLAB (MATLAB version R2010b Natick, Massachusetts: The MathWorks Inc.)

Centroid size was measured as the square root of the summed squared distances of 14 landmarks from the center of the wing (Fig. 1). Interocular distance was measured from eye edge to eye edge along the anterior edge of the posterior ocelli and parallel to the base of the head.

### Quantitative genetic analysis

All analyses were performed in R Studio using the R statistical language version 2.15 (http://www.R-project.org). PCA was performed on data of individual flies using the package *FactoMineR*. Allometric coefficients (b) were determined for each line and sex from the model log(y) = log(a) + b * log(x), where y = CS and x = IOD or TL, using the *lm()* function in the *stats* package. 95% confidence intervals for the parameter b were computed using the *confint()* function in the stats package. The total phenotypic variance in the control experiment was partitioned using the mixed model *Y = S + L + SxL + R(L) + ε*, where *S* is the fixed effect of sex, *L* is the random effect of line (genotype), *SxL* is the random effect of line by sex interaction, *R* is the random effect of replicate and *ε* is the within line variance. The brackets represent that replicate is nested within line. The total phenotypic variance in the dataset was partitioned using the mixed model *Y* = *S* + *L(F)* + *SxL(F)* + *F* + *ε*, where *S* is the fixed effect of sex, *L* is the random effect of line (genotype), *SxL* is the random effect of line by sex interaction, *F* is the random effect of food batch and *ε* is the within line variance. The random effects of line and line by sex are nested within food batch, as each line was raised only on one of the four food batches. Models of this form were fitted using the *lmer()* function in the *lme4* package in R. We also ran reduced models separately for males and females. The *rand()* function in the *lmerTest* package was used to assess significance of the random effects terms in the dataset.

Relative contributions of the variance components to total phenotypic variance 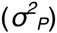 was calculated as 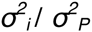 where 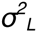 represents any of 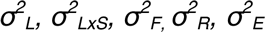, and 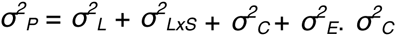 stands for 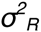 in the control dataset and for 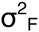 in the analysis of the GWAS dataset. 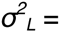 = variance due to genotype, 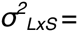 = variance due to genotype by sex interactions, 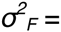 = variance due to food, 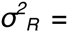 = variance due to replicate and 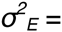 = residual (intra-line) variance. The broad sense heritability for each trait was estimated as 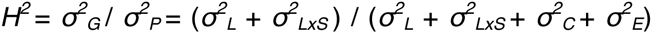. The cross-sex genetic correlation was calculated as 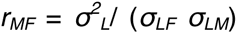 where 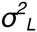 is the variance among lines from the analysis pooled across sexes, σ_*LF*_ and, σ_*LM*_ are, respectively, the square roots of the among line variance from the reduced models of females and males. Similarly, cross-trait genetic correlations were calculated as 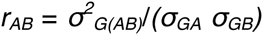 where 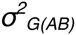 is the genetic covariance between traits A and B, and σ_*GA*_ and σ_*GB*_ are the square roots of the genetic variance for traits A and B, respectively. The phenotypic correlation between sexes was determined using the *cor()* function with method = “spearman” in R.

### Phenotypes for GWAS

We found a large effect of food batch on CS, and inversions *In(2L)t* and *In(3R)Mo* were associated with IOD and to a lesser extent CS. We modeled these covariates using a mixed model. The food batch was modeled by a random effect and the rearrangements were coded as (0,1,2) depending on whether both, one or no inversion was present in the homozygous state. We did not observe correlation between *Wolbachia* infection status and any trait and thus did not include this as a covariate in the model. Specifically, the models used were: *CS*_*raw*_ = *α* + *X*_*1*_*β*_*1*_ + *X*_*2*_*β*_*2*_ + *Fu* + *ε*, where *X*_*1*_ refers to the sex covariate, *X*_*2*_ refers to the inversion covariate, 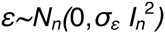 with *n* being the number of lines, 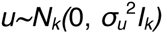 with *k* being the number of food batches and *Fu* an (*n,k*)-indicator matrix, associating each line to its respective food batch. The GWA analyses were performed using the estimated residual of this model (CS = *ε*).

To find loci that specifically affected variation in wing size unrelated to the overall body size variation we constructed an additional phenotype (rCS) that had the effect of IOD on CS removed via regression: *CS*_*raw*_ = *α* + *IOD* + *X*_*1*_*β*_*1*_ + *Fu* + *ε*, where *X*_*1*_ and *Fu* refer again to the sex-effect and the food batch effect. We did not model the inversions because the residuals of this model were not correlated with the inversions. The residuals *ε* from this regression were used as relative size phenotypes.

### Association analyses

We performed GWA analyses using male and female line means. Genotypes for 143 of the 149 lines were obtained from the DGRP Freeze 2 website (http://dgrp2.gnets.ncsu.edu). Only SNPs that were missing in a maximum of ten lines and occurred in at least ten lines (7% of the measured lines, 1,319,937 SNPs in total) were used. GWA was performed using FaST-LMM (53) for separate sexes. This association method uses a linear mixed model to capture confounders such as population structure and cryptic relatedness. Association results were visualized using the *manhattan()* function in the R package *qqman* (93). To determine correlation between SNPs for a given phenotype we extracted the genotype of the top *n* SNPs (*p*<10^-05^) and calculated the correlation between genotypes at these loci across all DGRP lines used in the GWA analyses. We used the FaST-LMM SNP *p*-values to apply the sum of chi-squares VEGAS method (56) to calculate gene wise statistics. Gene boundaries were defined using annotation from popDrowser (http://popdrowser.uab.cat/gb2/gbrowse/dgrp/), but we included also SNPs lying within 1,000 bp up- or downstream of these margins. The correlation matrix was calculated from the genotypes themselves.

### GO annotation and interaction enrichment

To determine enrichment of functional classes, annotate genes with functions and curated interactions among our candidate genes, we used the functional annotation and protein interaction enrichment tools from STRING (55).

### RNAi validation

SNPs with an association *p*-value <10^-05^ lying in a gene region or ± 1 kb from a gene were mapped to that gene. From the gene based VEGAS analysis, we chose the top 20 genes from each list as candidates. RNAi lines for a subset of candidate genes for each wing phenotype were ordered from VDRC (94). For one gene, *chinmo*, there was no appropriate line available from VDRC and we instead tested two Bloomington lines (26777 (y[1] v[1]; P{y[+t7.7] v[+t1.8]=TRiP.JF02341}attP2) and 33638 (y[1] v[1]; P{y[+t7.7] v[+t1.8]=TRiP.HMS00036}attP2/TM3, Sb[1]), indicated in S7 Table with (BL)). For the random control knockdowns we tested a set of 24 genes that did not contain a significant SNP in or within 1 kb of their transcribed region. We did the random knockdowns only in females to more effectively assess more genes for the same labor. As we wanted to address the controls like an additional phenotype (random) we chose a number of genes comparable to the numbers of candidates for other phenotypes. We chose females because we generally had more candidate genes in females than in males. All RNAi lines used are listed in S7 Table. For wing size candidates, validation was performed by crossing males of the respective RNAi line to virgin females carrying the *GAL4* transcriptional activator under the control of the *nubbin* (*nub*) promoter. The VDRC line containing a *UAS*-RNAi construct against the *CG1315* (GD library, transformant ID 47097) gene served as a negative control for the knockdowns. We decided to use this line as reference because it was in the same background as most of our tester lines, an essential factor to consider when assessing genes that presumably only have a small effect on size upon knockdown. The *CG1315* knockdown had never shown an effect in any setting and it allowed us to evaluate unspecific effects of RNAi knockdown on wing size. Prior to the experiment, driver lines were bred under controlled density to eliminate cross-generational effects of crowding on size. Wings were phenotyped as described above and wing area used as a phenotypic readout. Change in median wing area relative to the control was tested with a Wilcoxon rank sum test (function *wilcoxon.test()* in R) for each line and for separate sexes. If possible, 25 flies per cross and sex were phenotyped for statistical analysis, however sometimes the number of progeny was lower. The number of phenotyped flies per cross and sex is given in S7 Table. We used the *fisher.test()* function in R to determine if the proportion of validated genes was different among candidates and random lines, and the *wilcoxon.test()* function to test for a difference in median *p*-value between candidates and random lines. The comparison between candidate and random knockdowns was done for females exclusively as only this sex was measured for the random lines. Only genes not previously implied in wing development or growth control were included in the analysis, which excluded *chinmo*, *aPKC*, *tws* and *Ilp8* from the candidates and *EloA* and *spz5* from the random list.

### Epistatic analyses

We explored epistatic interactions between SNPs lying within and 1 kb around genes that were previously found to be involved in growth or wing development in *Drosophila* against all DGRP SNPs with missingness <11 and present in at least 10% of the lines. We compiled a list of SNPs within and 1 kb up- or down-stream of genes that were previously known to play a role in growth control (14,137 SNPs) or wing development (43,498 SNPs) and used these as focal SNPs (S8 Table). All phenotypes were normalized to follow a standard normal distribution for this analysis to make sure that no severely non-normal distributions occurred within any of the four marker classes per locus. We used FasT-Epistasis (57) for calculating interactions for all pairs between the focal SNPs and the set of all SNPs satisfying the above criteria (1,100,811 SNPs). Bonferroni corrected significance would thus require p<7.9×10^-13^. Interactions were visualized using Circos (95). To calculate significance for the overlap between genes found via epistasis and a given gene list, we first positionally indexed all n SNPs that were used in the epistasis analysis. We recorded the set of indices of SNPs with *p*<10^-09^) yielding set *K*: *K* = {k: *SNP*_*k*_ is an epistasis hit}. We then generated random samples. For random sample *j*, do:

For all elements in *K*, add a random integer *r*_*j*_ between *0* and *n-1*. Define new index as the modulo *n*: 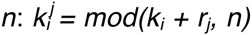, which yields 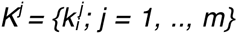. Given the shifted positions K^j^, we look up the SNP positions *P*_*Kj*_. For a given gene list, we record the number *x*_*j*_ of gene regions that overlap a position in *P*_*Kj*_. Let *x* be number of gene regions overlapping an epistasis hit. Our *p*-value estimate is then *P*_*approx*_ ≈ *1/m* ∑1_{xj ≥ x}_.

### Intergenic element enrichment analysis

We determined the number of SNPs from each GWA candidate list and the overall number of SNPs that located within modENCODE (65) elements annotated with Histone 3 lysine 4 monomethylation (H3K4Me1) or Histone 3 lysine 27 acetylation (H3K27Ac) or lincRNA loci. For the H3K4Me1/H3K27Ac enrichments we restricted ourselves to three developmental stages (L2, L3, pupae), which we considered to be the most relevant interval for gene activity affecting growth of imaginal discs. We obtained a table with lincRNAs in the *Drosophila* genome from the study of Young *et al.* (66) and searched for enrichment of SNPs located in those lincRNA loci. Enrichment was tested using a hypergeometric test (function *phyper()*) in R).

### BLAST alignment

We downloaded the sequence of the region 10 kb upstream of the annotated transcription start site of the *expanded* locus (*2L*: 421227..431227) from FlyBase (96), as well as the sequence of the same relative region for seven of the twelve *Drosophila* species (97), which contained the ortholog of the *expanded* gene in the same orientation in the genome. We performed multiple sequence alignment using the discontiguous megablast option on NCBI BLAST (69).

### Annotation with human orthologs

We combined candidate genes from GWA analyses of all phenotypes and searched for orthologs in humans using DIOPT-DIST (70). Enrichment of GWA candidates for genes with human orthologs associated with height (10) was determined with a hypergeometric test (function *phyper()* in R). We determined *Drosophila* orthologs of gene annotations of all associated SNPs (total 697), resulting in 374 ortholog pairs supported by at least 3 prediction tools, and searched for overlap of these orthologs with the 62 of our GWAS candidate genes that had a human ortholog supported by at least 3 prediction tools, which resulted in 12 matches. Of those, only five matches were supported by three or more prediction tools (score>=3) and we used only those for enrichment calculation. The total number of *Drosophila*-Human ortholog relationships (= 28,605) served as background (70).

## ACKNOWLEDGEMENTS

We thank Anna Troller, Benjamin Schlager and Anni Strässle for manual support during experiments. Stocks obtained from the Bloomington *Drosophila* Stock Center (NIH P40OD018537) and from the Vienna *Drosophila* RNAi Center were used in this study.

## AUTHOR CONTRIBUTIONS

EH, TM and SB conceived the approach. SV designed and performed experiments and SV and DL performed statistical analyses. SV wrote the manuscript and SV, DL, EH, SB and TM edited the manuscript.

## SUPPORTING INFORMATION

**S1 Fig.**
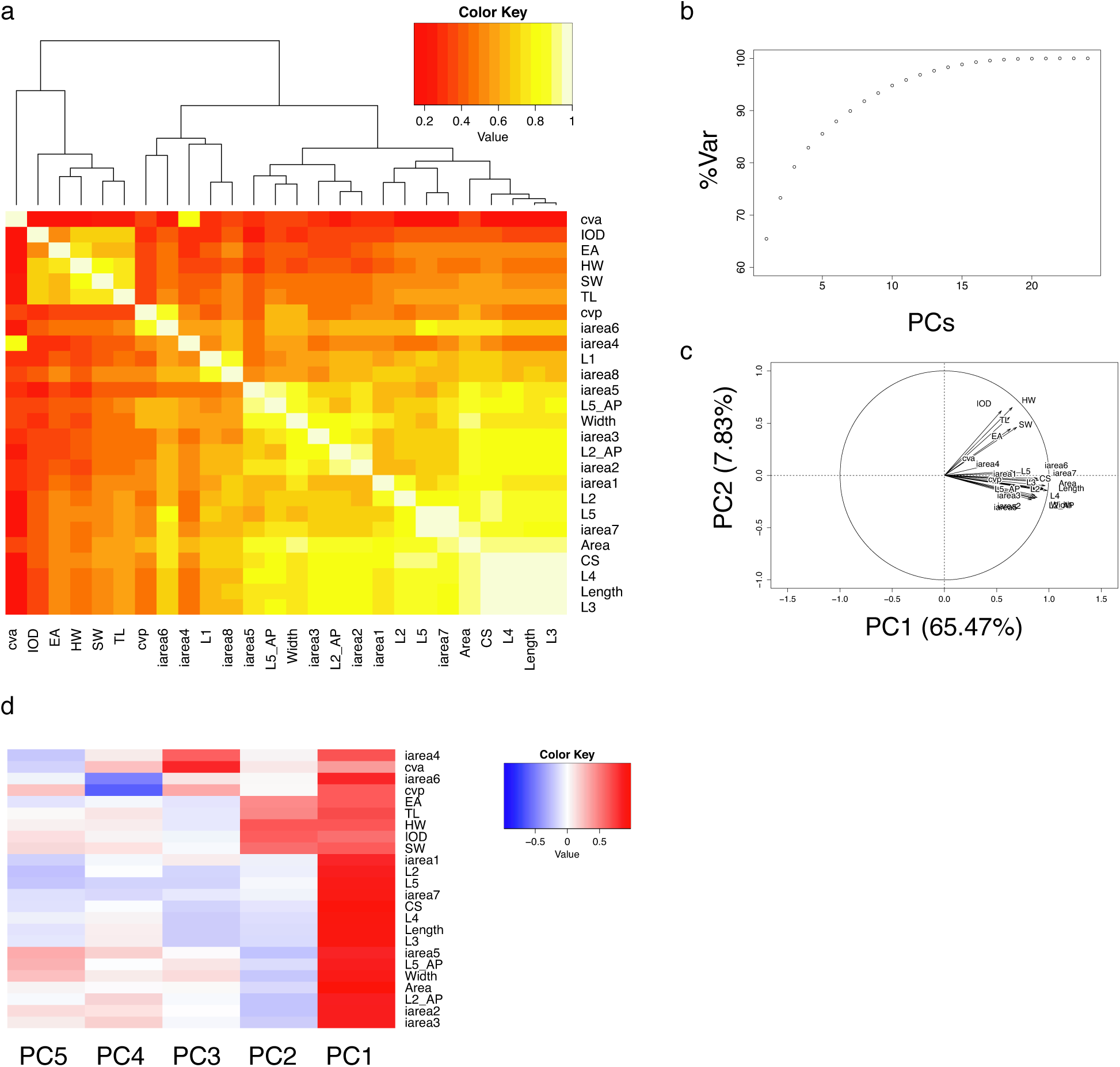
Analysis of the male dataset. a) Genetic correlation between morphometric traits in males. The two modules of higher correlation observed in females are still visible (bright yellow in the upper left and lower right corners) but the overall clustering is more influenced by the more inaccurately measured smaller veins and areas. b) Cumulative variance explained in male data by increasing number of principal components. As in the female dataset, the first two PCs explain almost 75% of the variance in the data. c) Factor map for the variables. PCs 1 and 2 split the data into two groups. d) Correlation between PCs and traits. PC1 reflects a general size component and PC2 is highly correlated with head/thorax traits, effectively splitting the data in two groups.

**S2 Fig.**
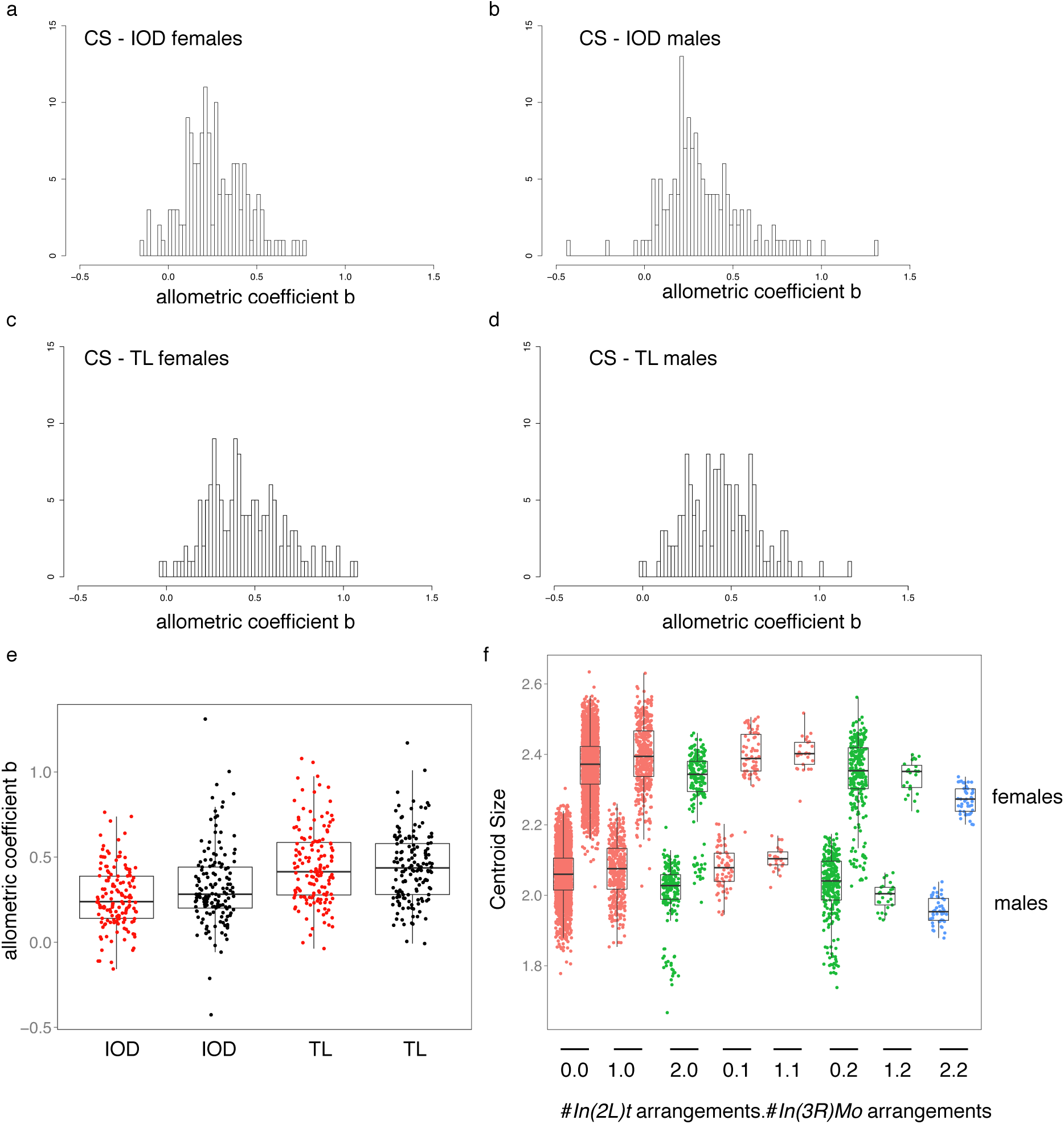
Allometry and inversions. Histograms of the estimates for the allometric coefficient *b* for the relationship between CS and IOD in females (a), in males (b) and between CS and TL in females (c) and males(d). e) Boxplot and individual datapoints of the data in a-d. Red = females and black = males. 95% confidence intervals for *b* (S1 Table) are very broad for some lines due to few datapoints used for fitting, so these are just very rough estimates for the allometric relationship. Nevertheless there is variation among lines for all evaluated relationships. f) The effect of cosmopolitan inversions on wing size. Lines are plotted according to the number of homozygous inversion arrangements they have: 0 (red) = neither *In(2L)t* nor *In(3R)Mo* present, 1 (green) = homozygous for either *In(2L)t* or *In(3R)Mo*, 2 (blue)=homozygous for both *In(2L)t* and *In(3R)Mo*. Datapoints are individual flies.

**S3 Fig.**
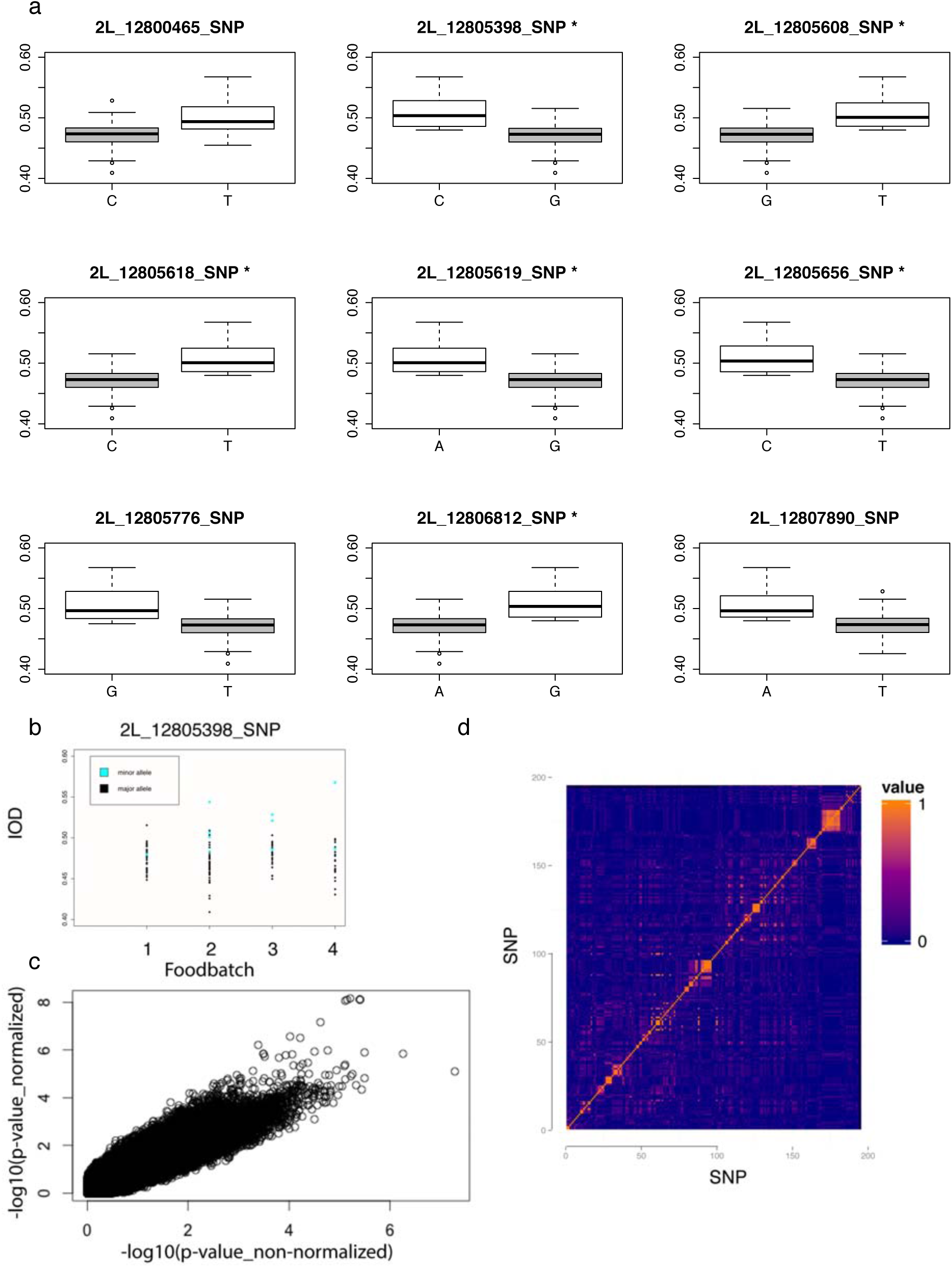
The minor and major haplotype of genome-wide significant SNPs show differential association with female IOD. a) The minor allele haplotype of the genome-wide significant cluster is associated with an increased IOD in females. Boxplots of female IOD by genotype at the nine SNPs annotated to *kek1*. SNPs marked by a star pass Bonferroni correction. Grey=major allele, white = minor allele. b) Lines with the minor haplotype are distributed across all four foodbatches. Black dots = major allele, blued dots = minor allele. IOD by foodbatch is plotted for females for the most significant SNP. The distribution is the same for all other SNPs of the cluster as all minor alleles form a haplotype. c) Correlation between *p*-values from GWAS with normalized IOD (y-axis) and non-normalized iod (x-axis) in females. Axes are on the –log10 scale. d) Several blocks of higher LD are visible in the region 20kb upstream of *kek1*. Blue = no correlation, orange = complete correlation.

**S4 Fig.**
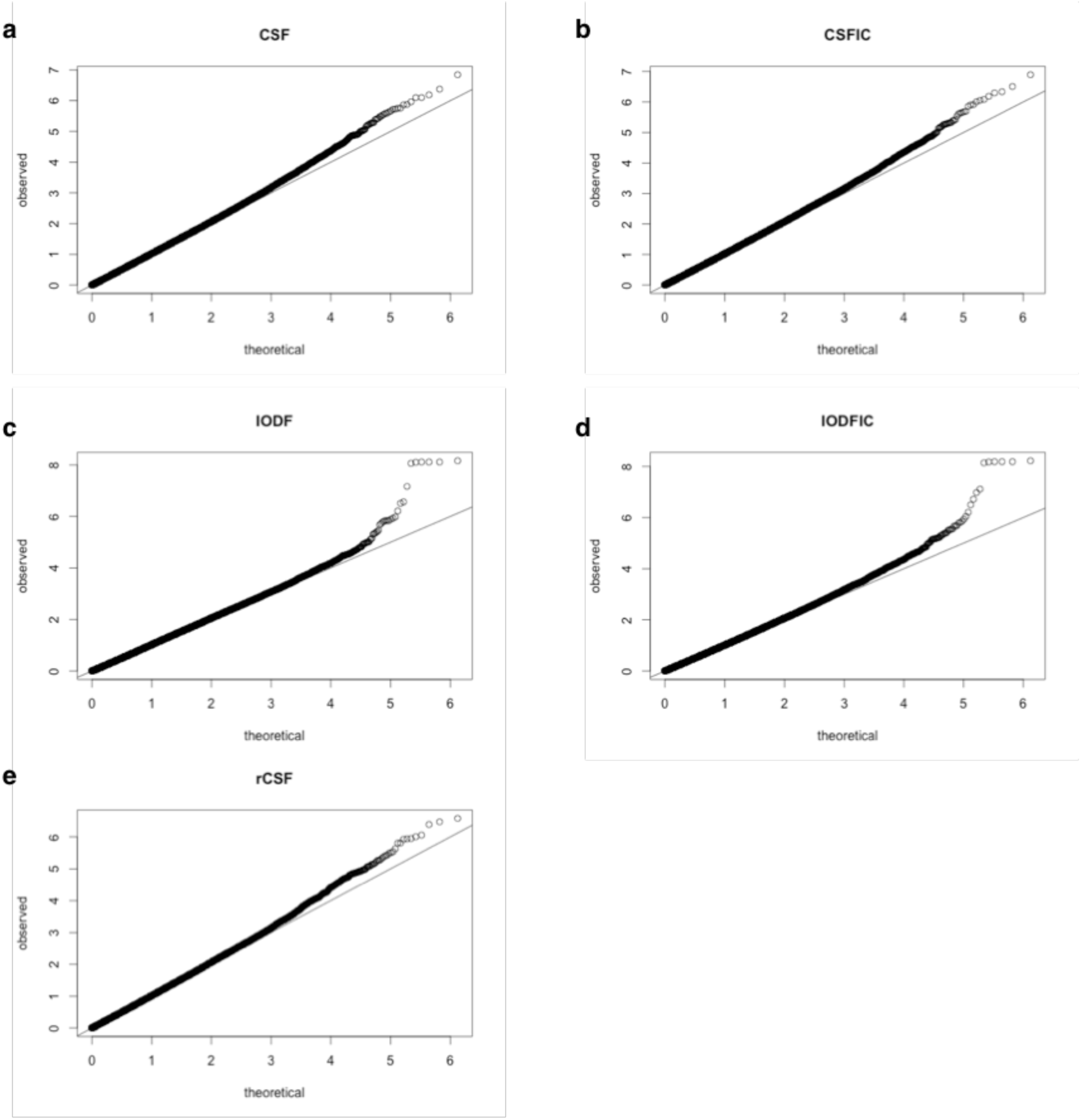
QQ-plots from GWA in females for all traits show a departure from uniformity of top associations. Observed association *p*-values are –log10 transformed (y-axis) and plotted against the –log10 transformed theoretically expected *p*-values under the assumption of no association (uniform distribution, x-axis). Centroid size (a), inversion corrected centroid size (b), interocular distance (c), inversion corrected interocular distance (d) and relative centroid size (e).

**S5 Fig.**
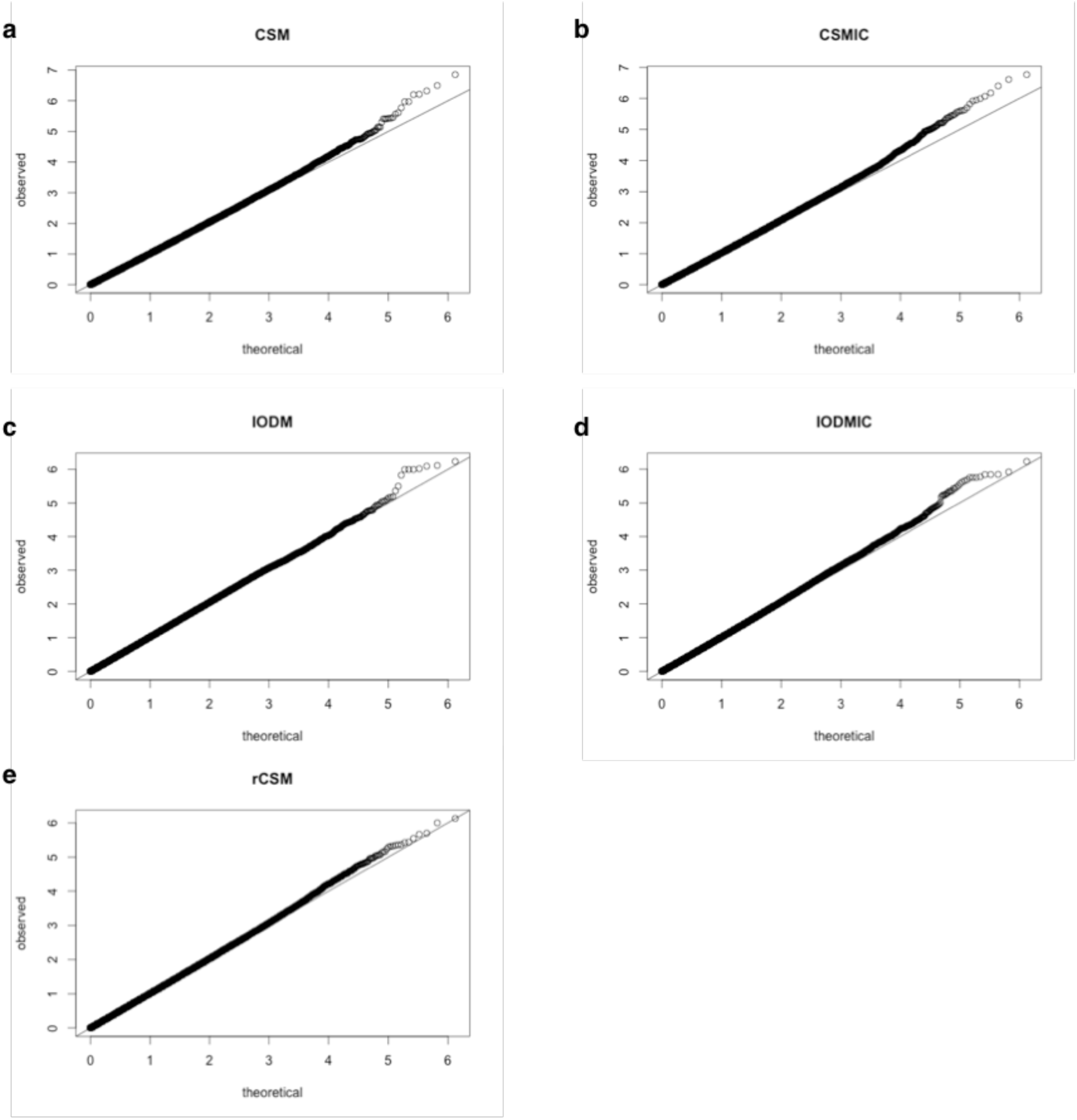
QQ-plots from GWAS in males for all traits show a departure from uniformity of top associations. Observed association *p*-values are –log10 transformed (y-axis) and plotted against the –log10 transformed theoretically expected *p*-values under the assumption of no association (uniform distribution, x-axis). Centroid size (a), inversion corrected centroid size (b), interocular distance (c), inversion corrected interocular distance (d) and relative centroid size (e).

**S6 Fig.**
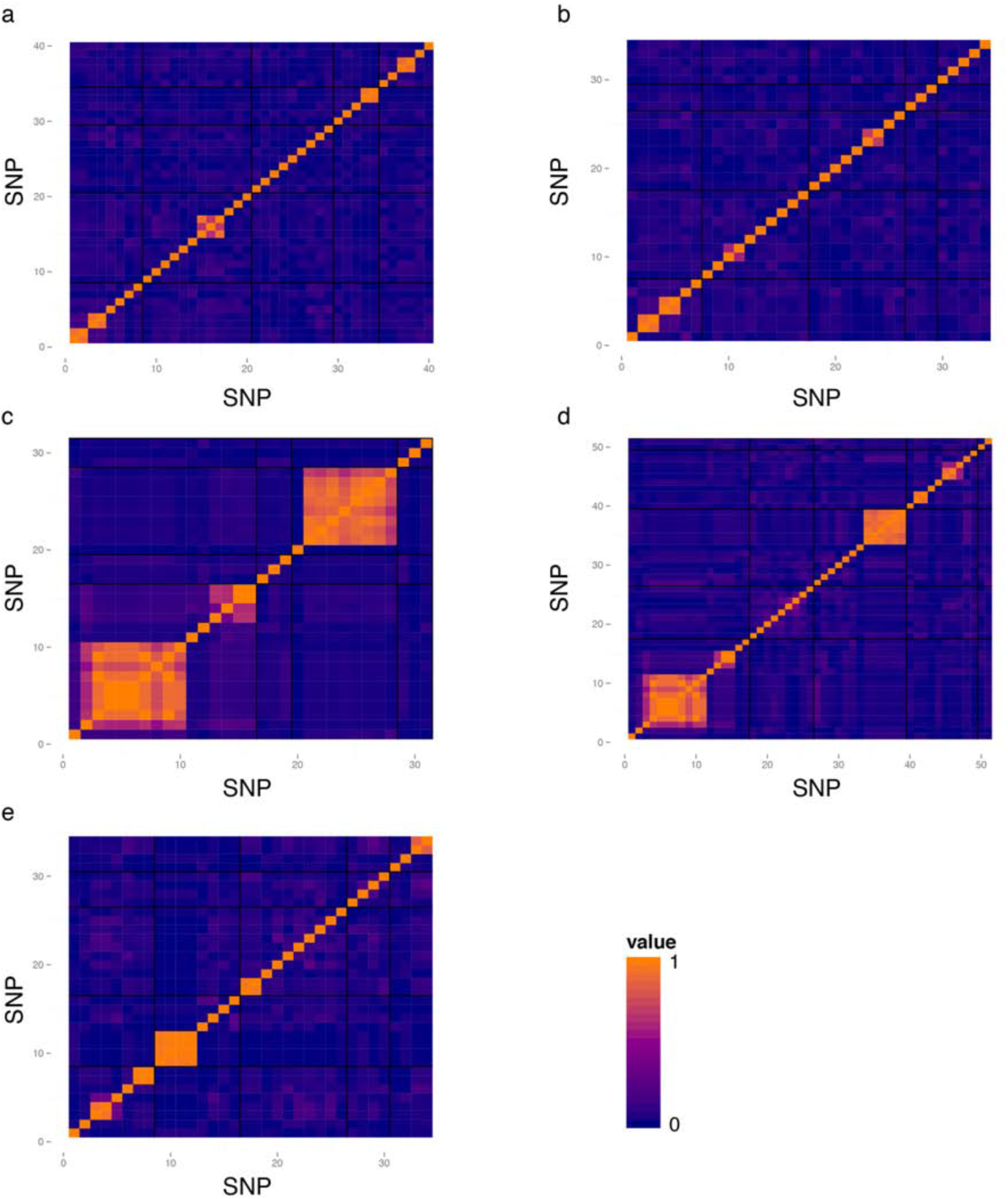
Correlation between associated (*p* <10^-05^) SNPs in females. The SNPs are ordered according to chromosome arm (*2L*, *2R*, *3L*, *3R*, *X*) and black dividers separate chromosomes. Within one chromosome arm SNPs are ordered according to their position on that chromosome with each tile representing one SNP. The color code is depicted on the right: orange = complete correlation (1) and blue = no correlation (0). Centroid size (a), inversion corrected centroid size (b), interocular distance (c), inversion corrected interocular distance (d) and relative centroid size (e).

**S7 Fig.**
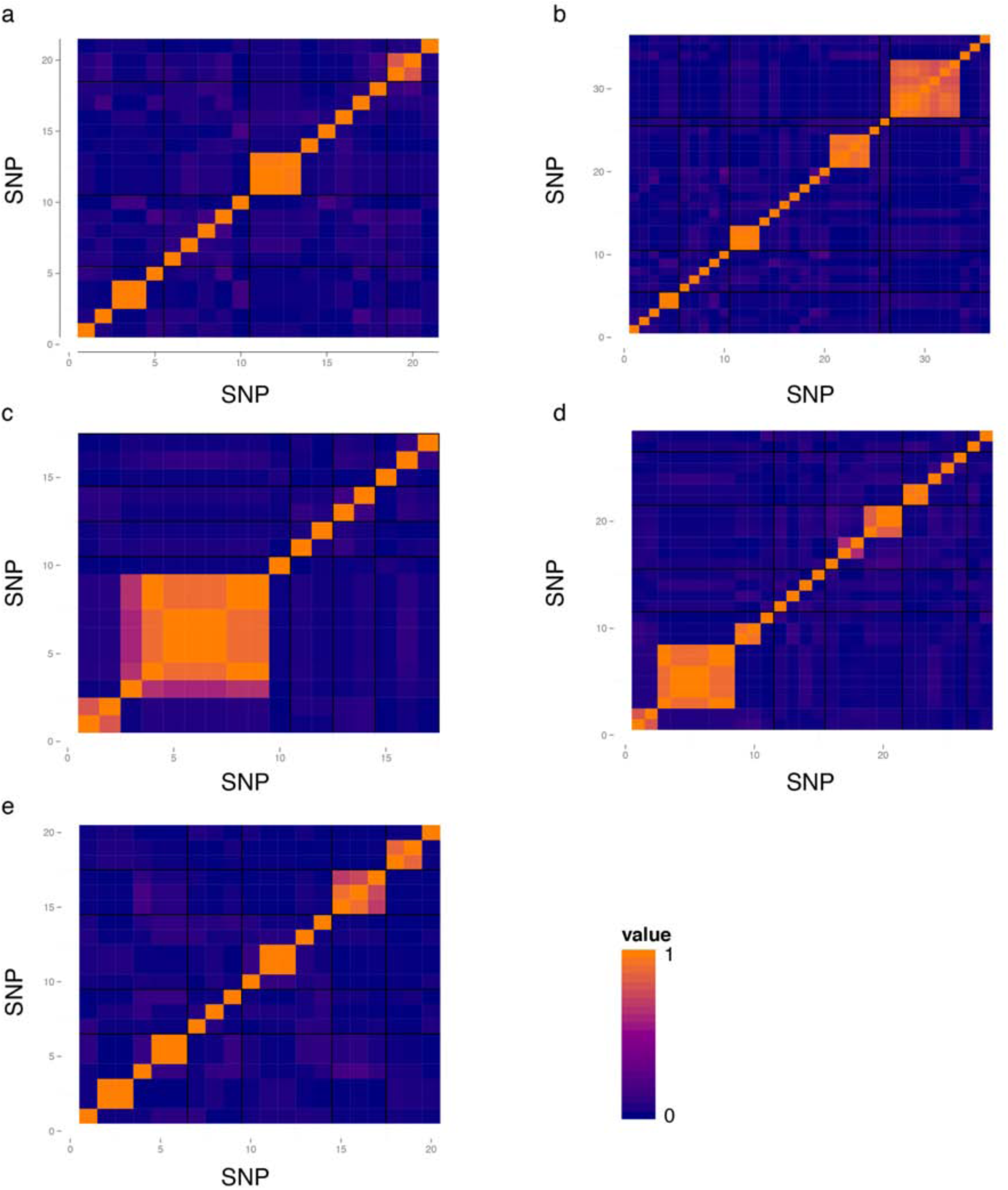
Correlation between associated (*p*<10^-05^) SNPs in males. The SNPs are ordered according to chromosome arm (*2L*, *2R*, *3L*, *3R*, *X*) and black dividers separate chromosomes. Within one chromosome arm SNPs are ordered according to their position on that chromosome with each tile representing one SNP. The color code is depicted on the right: orange = complete correlation (1) and blue = no correlation (0). Centroid size (a), inversion corrected centroid size (b), interocular distance (c), inversion corrected interocular distance (d) and relative centroid size (e).

**S8 Fig.**
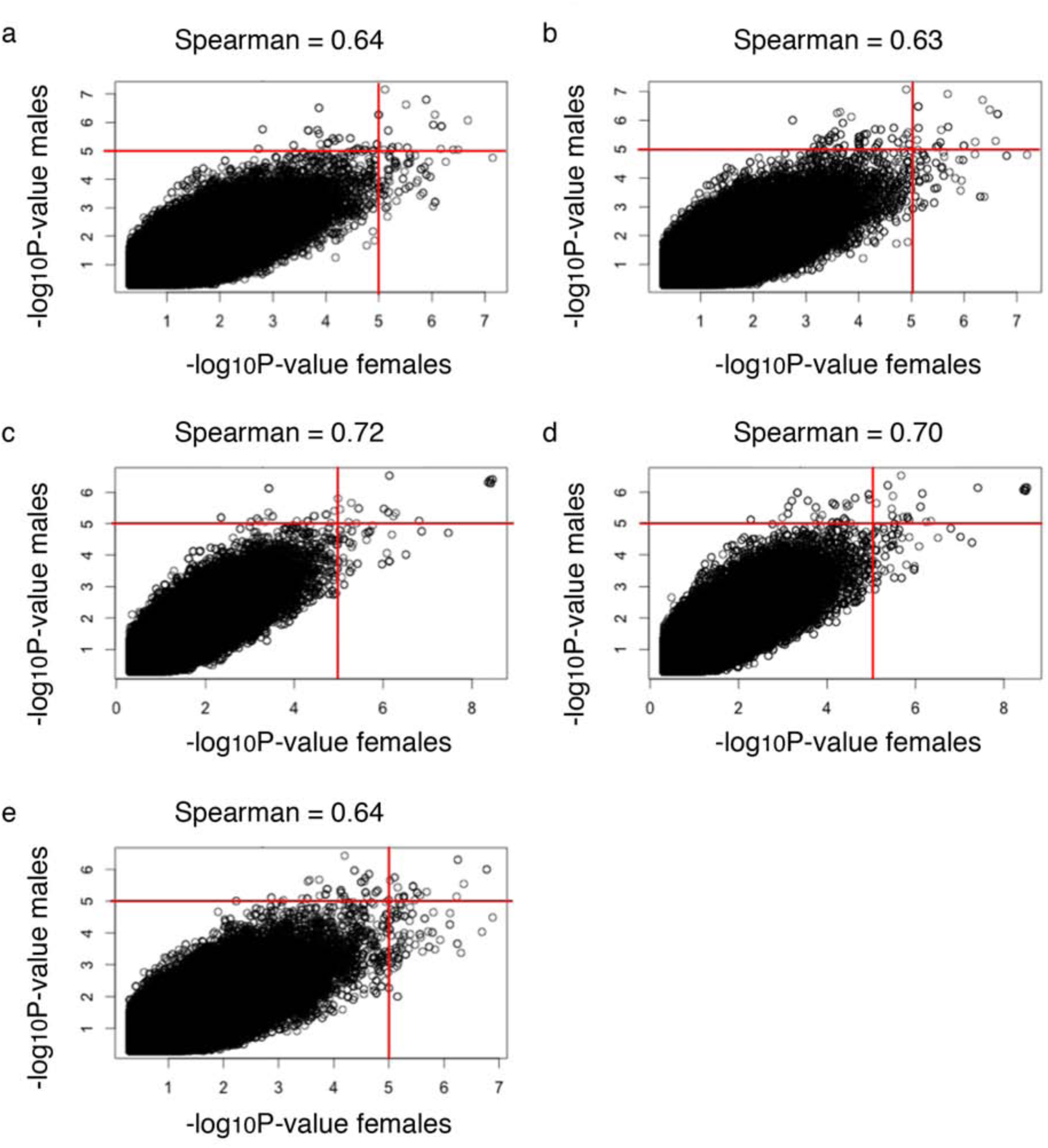
Correlation of SNP *p*-values between the sexes. SNP *p*-values in females (x-axis) are plotted against their respective p-values in males (y-axis). The Spearman rank correlation is given for each trait and the red lines denote the significance cutoff. a = CS, b = CS_IC_, c = IOD, d = IOD_IC_, e = rCS.

**S9 Fig.**
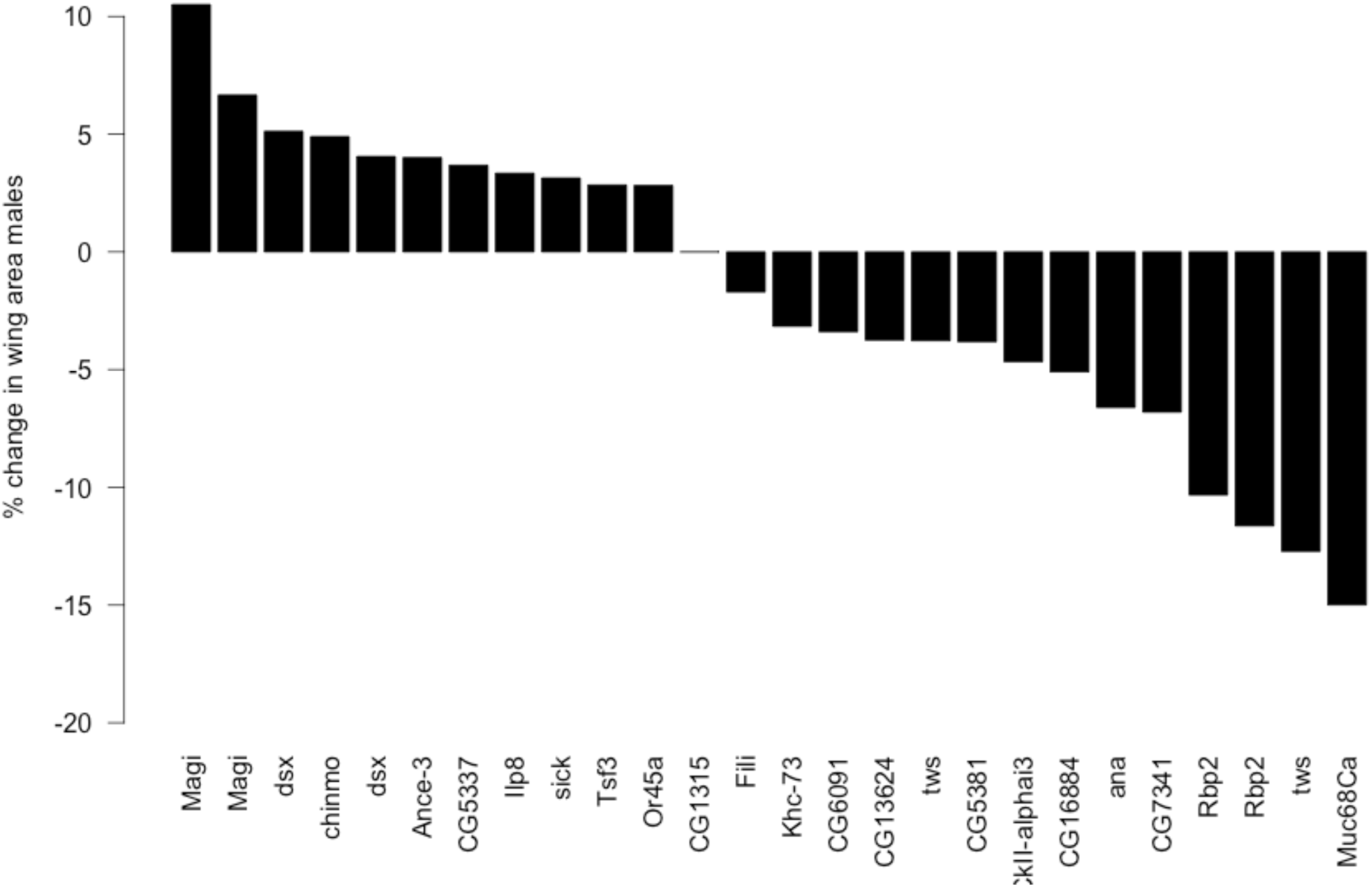
RNAi knockdown results males. Percent change in median wing area compared to *CG1315* RNAi upon wing-specific knockdown of the validated candidate genes in males. Only the lines yielding a significant wing size change (*p*<0.001, Wilcoxon rank sum test) are depicted. Median, 25^th^ and 75^th^ percentile for each are given in S7 Table.

**S10 Fig.**
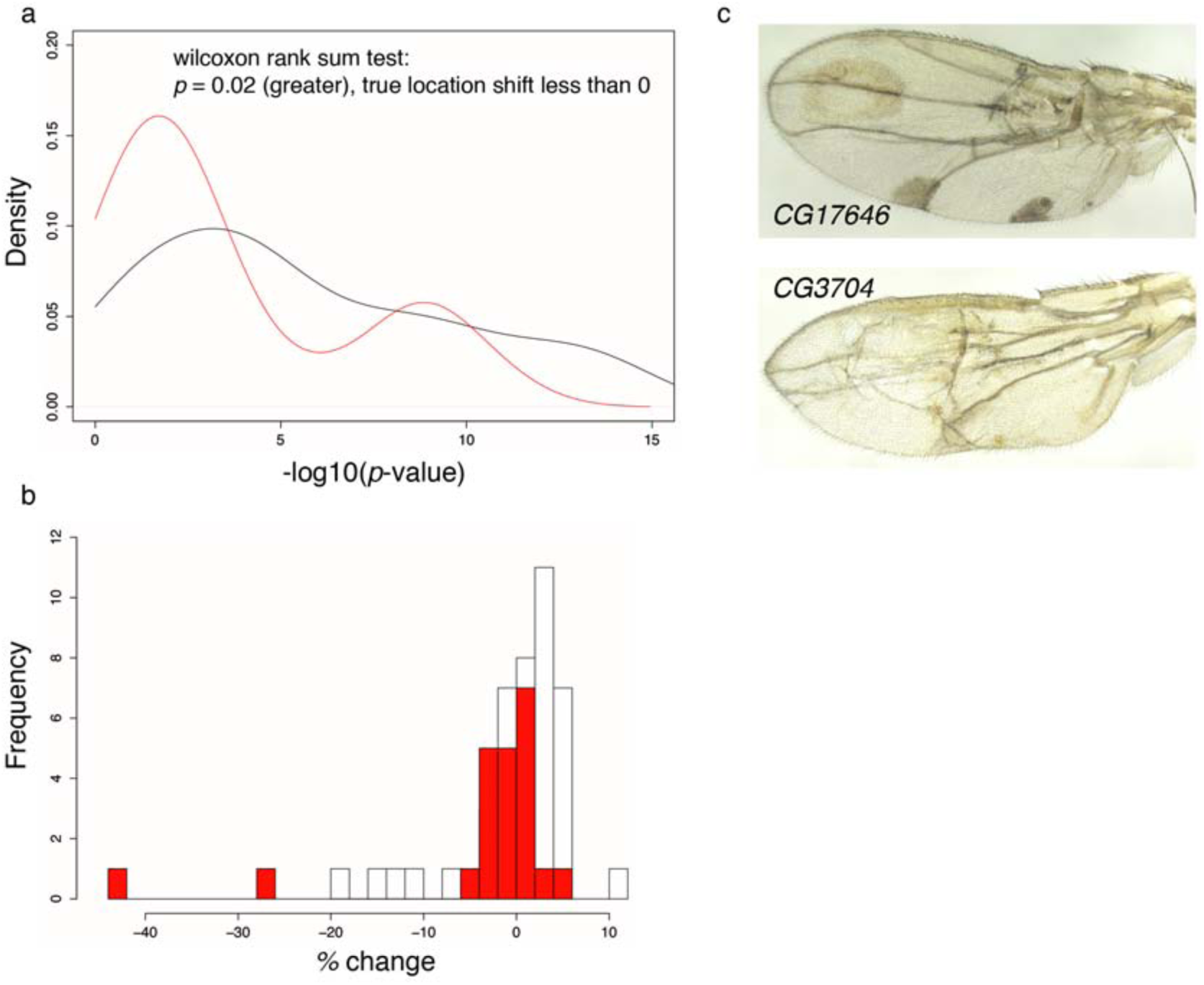
Comparison of *p*-values and effect sizes between candidate and control RNAi. a: -log10 transformed *p*-value densities of the candidate (black) and combined control (red) data sets. The two *p*-value distributions differ by a location shift that is not zero (i.e. are not the same); specifically, the – log10 transformed control *p*-value distribution (red) is shifted towards the left of the – log10 transformed candidate *p*-value distribution (black) (one sided Wilcoxon rank sum test *p* = 0.02). b: The distribution of candidate effect sizes (percent change in wing size upon knockdown) is shifted towards positive effect sizes (white boxes), whereas the control knockdown effect size distribution (red) is more centered on 0. The two exceptions at -28% (*CG17646*) and -42% (*CG3704*) are lines whose wings not only show a size reduction but also considerable morphological defects (c). *N* = 43 candidates (white), *N* = 22 control (red); only data from females was used for these analyses.

**S11 Fig.**
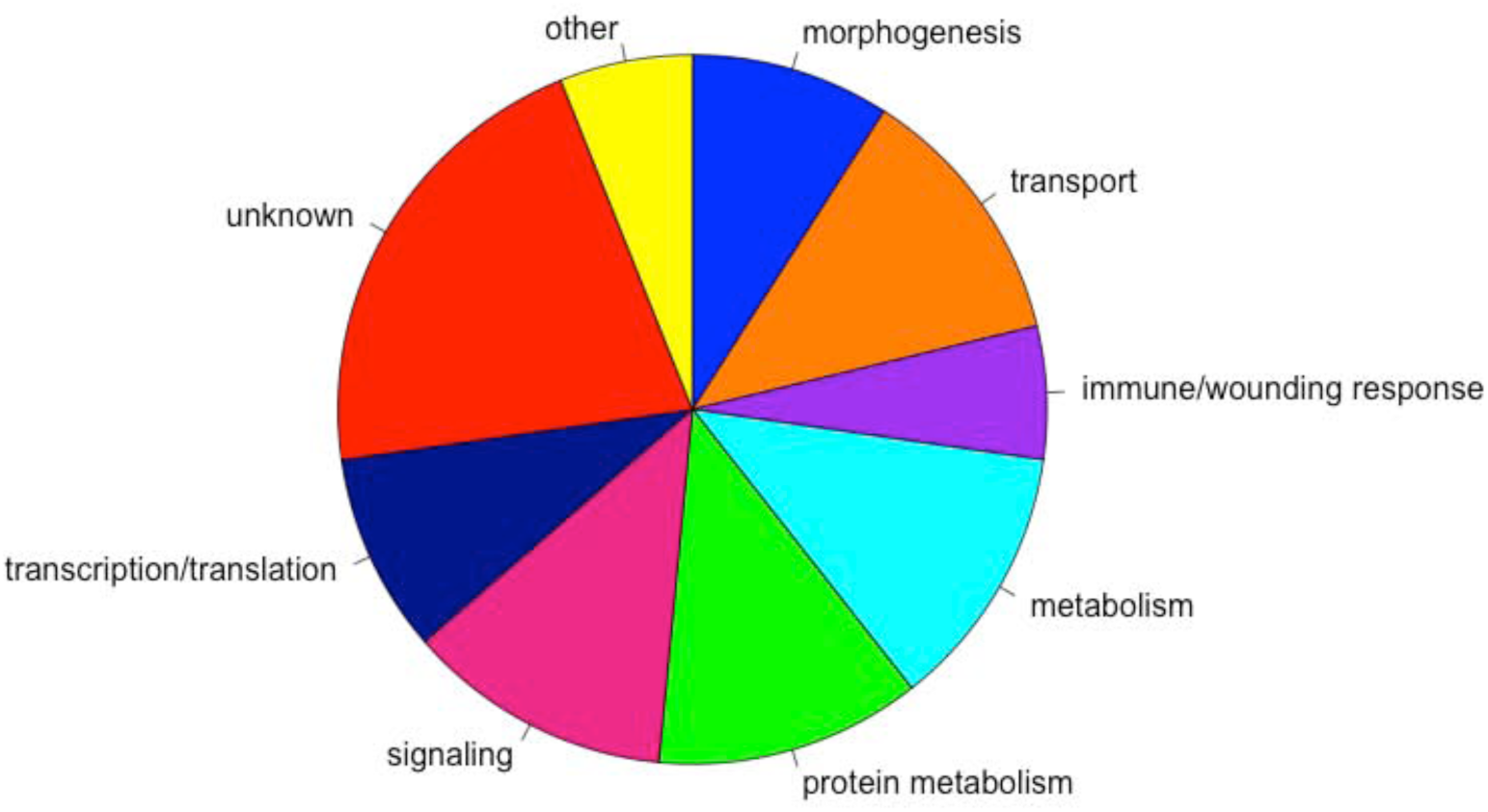
Functional annotation of the 33 validated candidate genes based on DAVID GO annotation.

**S1 Table. Quantitative genetic analysis.** Control experiment: Only 2% of total population variance in CS and IOD was due to flies coming from replicate vials, a negligible fraction compared to the 78% and 71% attributable to differences in genotype. This indicates that the standardized culture protocol sufficiently deals with confounding effects on size phenotypes. *N*=2000, 25 flies/sex of 10 replicates of four DGRP lines. Allometry: Allometric coefficients (b) calculated from the equation log(CS) = log(a) + b*log(trait) and their 95% confidence interval (CI) are given for the allometric relationships between CS and IOD and CS and TL for each line and sex. Though the CI varies substantially for some lines due to few data points used for fitting the models, the upper CI boundary is close to 0 for some (e.g. 28157 females (RAL228) for the CS-IOD relationship). QGA Dataset: Quantitative genetic analysis of CS and IOD in the dataset consisting of *N*=6978 flies from 149 DGRP lines. *p*_Sex_ = significance of fixed effect of sex, *p*_Line_ = significance of random effect of Line, *p*_Sex x Line_ = significance of random effect of Line by sex interaction, *p*_Replicate_ = significance of random effect of replicate, *p*_Food_ = significance of random effect of foodbatch. Estimated parameters are variance due to genotype (V_G_), genotype by sex interaction (V_GxS_), food (V_F_), replicate (V_R_) and intra-line variance (V_E_), as well as the cross-sex genetic (r_MF_) and phenotypic correlation between sexes. H^2^ = broad-sense heritability and V_P_ = total phenotypic variance. Phenotypic variation: Smallest and largest trait values and the percent difference are given per sex for each phenotype. Population means (Mean) and standard deviations (STD) are used to calculate the coefficient of variation (CV).

**S2 Table. Phenotypic data.** Raw data for all traits and line means (Mean), standard deviations (STD) and number of phenotyped flies (N) listed by sex for centroid size (CS) and interocular distance (IOD).

**S3 Table. GWAS results.** Nominally significant SNPs (*p*<10^-05^) from GWAS for Centroid size (CS), inversion modeled centroid size (CS_IC_), interocular distance (IOD), inversion modeled interocular distance (IOD_IC_) and relative centroid size (rCS) in both sexes.

**S4 Table. Between-sex and -phenotype overlap of nominally significant SNPs.** GWAS SNPs: Number of nominally significant SNPs (*p*<10^-05^) identified in each of the GWAS and percent overlap between sexes. SNPs F = number of SNPs nominally significant in females, SNPs M = number of SNPs nominally significant in males, Common MF = number of SNPs nominally significant in both sexes, % M in F = percent nominally significant male SNPs also nominally significant in females, % F in M = percent nominally significant female SNPs also nominally significant in males. Between sex overlap: SNP and gene level overlap between sexes for different thresholds. Percent overlap of SNPs (exact associated positions), annotated SNPs (including annotation of SNPs located in intergenic regions where the annotation is often not reliable, as the next genes are frequently more than 20kb away), and genes (only includes SNPs that locate within or 1kb around a gene). A consistent difference in associated loci in terms of the exact SNP that is associated (top block) is detectable between sexes, though the loci apparently are located more or less in a similar region (middle block). For SNPs in or close to genes, the differences between sexes are more pronounced: of the top 100 associated genes about 50% overlap, while the other 50% are private to one sex but this percentage increases with inclusion of more genes (bottom block, top 10,000 is >90% overlap, in total there are around 14,000 genes in these lists). Between phenotype overlap: Proportion of nominally significant SNPs that are shared between phenotypes.

**S5 Table. Enrichment analysis.** Enrichment: Enrichment *p*-values for SNPs localizing to different genomic regions. LincRNA: LincRNA loci that overlap significant SNPs and their expression during different developmental stages. LincRNA data from Young *et al.* (54).

**S6 Table. Top 20 genes for each trait identified by the VEGAS method.** Last column indicates the percent change in median wing size of genes tested by RNAi, and asterisks (***) indicate significant change (*p*<0.001).

**S7 Table. Validation results candidates and random genes.** The lines showing a significant (*p*<0.001, Wilcoxon rank sum test) change in median wing area upon knockdown are shaded in grey. Crosses in columns pigmentation, bristles, veins indicate a slightly abnormal corresponding phenotype. N= number of individuals tested, Median = median wing area, Q25 and Q75 = first and third quartile of wing area distribution. MAF 3 or MAF 5 in brackets means a SNP in or near this gene was found among the top associated genes in a GWAS for wing size with a lower MAF cut-off (SNPs present in min. 3% or 5% of lines). This gene was tested for wing size since a SNP in it showed association to body size with the used MAF cutoff and thus a corresponding RNAi line was available. Overview: Overview validation results. Number of SNPs that were tested and the number and percentage that were validated for each trait. We tested the known genes *chinmo*, *aPKC*, *tws* and *Ilp8* as positive controls, but did not include them in the calculation of these percentages. Fisher’s exact test: We performed a two-sided Fisher’s exact test to determine if the proportions of validated genes was different between candidates and random genes. The results are shown for different Wilcoxon test *p*-value validation thresholds. Only not previously known candidates and random lines were used (N_candidates_ = 43, N_random_ = 22)

**S8 Table. Epistasis results.** Focal genes: Genes previously implied in growth regulation or wing development that were used as focal genes for epistasis. Top interactions (*p*<10^-09^) for female absolute inversion corrected wing size (CSF_IC_), male absolute inversion corrected wing size (CSM_IC_), female absolute inversion corrected body size (IODF_IC_), male absolute inversion corrected body size (IODM_IC_), female relative wing size (rCSF) and male relative wing size (rCSM). X is the interactor locus and Y the focal (= previously known) locus.

**S9 Table. Multiple sequence alignment (MSA) results.** *Expanded* orthologs: Genomic location of *ex* orthologs in 12 *Drosophila* species (82). Name of the ortholog, its genomic location and orientation are shown. For the MSA we only used species that contained the gene in the same orientation as *D. melanogaster* (grey). MSA details: Details of MSA of the 10kb region upstream of *D. melanogaster ex* gene. Sequence = genomic region in each species used in the MSA, Identity = percent identical nucleotides, Aligned Length = length of alignment, Query Cover = percent of the query sequence (*D. melanogaster* sequence) aligned to the sequence in the respective species, E-value = significance of alignment (expected number of high scoring pairs with score at least as high as the score of the current alignment), Score = strength of alignment.

**S10 Table. Human orthologs of putative and validated *Drosophila* growth regulators and their association to human complex traits.**

**S11 Table. GWAS/epistasis candidates reported by other studies.** Candidates found as suppressors or enhancers of major growth pathways by Schertel *et al.* (64). Candidates associated with nutritional indices in the study of Unckless *et al.* (65)

## REFERENCES

1. Oldham S, Bohni R, Stocker H, Brogiolo W, Hafen E. Genetic control of size in Drosophila. Phil. Trans. R. Soc. B: Biological Sciences 2000; 355: 945–952.

2. Johnston LA, Gallant P. Control of growth and organ size in Drosophila. Bioessays 2002; 24: 54–64.

3. Mirth CK, Riddiford LM. Size assessment and growth control: how adult size is determined in insects. Bioessays 2007; 29: 344–355.

4. Shingleton AW. The regulation of organ size in Drosophila: physiology, plasticity, patterning and physical force. Organogenesis 2010; 6: 76–87.

5. Oldham S, Hafen E. Insulin/IGF and target of rapamycin signaling: a TOR de force in growth control. Trends Cell Biol.2003; 13: 79–85.

6. Pan D. Hippo signaling in organ size control. Genes Dev. 2007; 21: 886–897.

7. Tumaneng K, Russell RC, Guan KL. Organ size control by Hippo and TOR pathways. Curr. Biol. 2012; 22: R368–R379.

8. Gockel J, Robinson SJW, Kennington WJ, Goldstein DB, Partridge L. Quantitative genetic analysis of natural variation in body size in Drosophila melanogaster. Heredity 2002; 89: 145–153.

9. Lango Allen H, Estrada K, Lettre G, Berndt SI, Weedon MN, Rivadeneira F, et al. Hundreds of variants clustered in genomic loci and biological pathways affect human height. Nature 2010;467: 832–838 (2010).

10. Wood AR, Esko T, Yang J, Vedantam S, Pers TH, Gustafsson S, et al. Defining the role of common variation in the genomic and biological architecture of adult human height. Nat. Genet. 2014;46: 1173–1186.

11. Falconer DS, Mackay TFC. Introduction to Quantitative Genetics. 4th ed. Harlow, Essex, UK: Longmans Green; 1996.

12. Lynch M, Walsh B. Genetics and Analysis of Quantitative Traits. Sunderland MA, USA: Sinauer Associates, Inc; 1998.

13. Robertson FW, Reeve E. Studies in quantitative inheritance I. The effects of selection of wing and thorax length in Drosophila melanogaster. J. Genet. 1952; 50: 414–448.

14. Partridge L, Langelan R, Fowler K, Zwaan B, French V. Correlated responses to selection on body size in Drosophila melanogaster. Genetics Research 1999; 74: 43–54.

15. Trotta V, Calboli FCF, Ziosi M, Cavicchi S. Fitness variation in response to artificial selection for reduced cell area, cell number and wing area in natural populations of Drosophila melanogaster. BMC Evol. Biol.2007; 7: Suppl 2, S10.

16. Turner TL, Stewart AD, Fields AT, Rice WR, Tarone AM. Population-Based Resequencing of Experimentally Evolved Populations Reveals the Genetic Basis of Body Size Variation in Drosophila melanogaster. PLoS Genet. 2011; 7: e1001336.

17. Gockel J, Robinson SJW, Kennington WJ, Goldstein DB, Partridge L. Quantitative genetic analysis of natural variation in body size in Drosophila melanogaster. Heredity 2002; 89: 145–153.

18. Calboli FCF, Kennington WJ, Partridge L. QTL mapping reveals a striking coincidence in the positions of genomic regions associated with adaptive variation in body size in parallel clines of Drosophila melanogaster on different continents. Evolution 2003; 57: 2653–2658.

19. Rako L, Anderson AR, Sgro CM, Stocker AJ, Hoffmann AA. The association between inversion In(3R)Payne and clinally varying traits in Drosophila melanogaster. Genetica 2006; 128: 373–384.

20. Kennington WJ, Hoffmann AA, Partridge L. Mapping regions within cosmopolitan inversion In(3R)Payne associated with natural variation in body size in Drosophila melanogaster. Genetics 2007; 177: 549–556.

21. Weeks AR, McKechnie SW, Hoffmann AA. Dissecting adaptive clinal variation: markers, inversions and size/stress associations in Drosophila melanogaster from a central field population. Ecology Letters 2002; 5: 756–763.

22. DeJong G, Bochdanovits Z. Latitudinal clines in Drosophila melanogaster: body size, allozyme frequencies, inversion frequencies, and the insulin-signalling pathway. J. Genet. 2003; 82: 207–223.

23. McKechnie SW, Blacket MJ, Song SV, Rako L, Carroll X, Johnson TK, Jensen LT, Lee SF, Wee CW, Hoffmann AA. A clinally varying promoter polymorphism associated with adaptive variation in wing size in Drosophila. Mol. Ecol. 2010; 19: 775–784.

24. Paaby AB, Bergland AO, Behrman EL, Schmidt PS. A highly pleiotropic amino acid polymorphism in the Drosophilainsulin receptor contributes to life-history adaptation. Evolution 2014; 68: 3395–3409.

25. Womack JE, Jang HJ, Lee MO. Genomics of complex traits. Ann. N. Y. Acad. Sci. 2012; 1271: 33–36.

26. Korte A, Farlow A. The advantages and limitations of trait analysis with GWAS-a review. Plant Methods 2013; 9: p. 29..

27. Hirschhorn JN, Daly MJ. Genome-wide association studies for common diseases and complex traits. Nat. Rev. Genet. 2005; 6: 95–108.

28. McCarthy MI, Abecasis GR, Cardon LR, Goldstein DB, Little J, Ioannidis JPA, et al. Genome-wide association studies for complex traits: consensus, uncertainty and challenges. Nat. Rev. Genet. 2008; 9: 356–369.

29. Bergelson J, Roux F. Towards identifying genes underlying ecologically relevant traits in Arabidopsis thaliana. Nat. Rev. Genet. 2010; 11: 867–879.

30. Meijón M, Satbhai SB, Tsuchimatsu T, Busch W. Genome-wide association study using cellular traits identifies a new regulator of root development in Arabidopsis. Nat. Genet.2013; 46: 77–81.

31. Jumbo-Lucioni P, Ayroles JF, Chambers M, Jordan KW, Leips J, Mackay TFC, et al. Systems genetics analysis of body weight and energy metabolism traits in Drosophila melanogaster. BMC Genomics 2010; 11: 297.

32. Jumbo-Lucioni P, Bu S, Harbison ST, Slaughter JC, Mackay TFC, Moellering DR, et al. Nuclear genomic control of naturally occurring variation in mitochondrial function in Drosophila melanogaster. BMC Genomics 2012; 13: p. 659.

33. Swarup S, Huang W, Mackay TFC, Anholt RRH. Analysis of natural variation reveals neurogenetic networks for Drosophila olfactory behavior. Proc. Natl. Acad. Sci. USA 2013; 110: 1017–1022.

34. Flint J, Eskin E. Genome-wide association studies in mice. Nat. Rev. Genet. 2012; 13: 807–817.

35. Huang X, Zhao Y, Wei X, Li C, Wang A, Zhao Q, et al. Genome-wide association study of flowering time and grain yield traits in a worldwide collection of rice germplasm. Nat. Genet. 2011; 44: 32–39.

36. Lipka AE, Gore MA, Magallanes-Lundback M, Mesberg A, Lin H, Tiede T, et al. Genome-wide association study and pathway-level analysis of tocochromanol levels in maize grain. G3 (Bethesda) 2013; 3: 1287–1299.

37. García-Gámez E, Gutiérrez-Gil B, Sahana G, Sànchez JP, Bayón Y, Arranz JJ. GWA Analysis for Milk Production Traits in Dairy Sheep and Genetic Support for a QTN Influencing Milk Protein Percentage in the LALBA Gene. PLoS ONE 2012; 7: e47782.

38. Makvandi-Nejad S, Hoffman GE, Allen JJ, Chu E, Gu E, Chandler AM, et al. Four Loci Explain 83% of Size Variation in the Horse. PLoS ONE 2012; 7: e39929.

39. Maxa J, Neuditschko M, Russ I, Förster M, Medugorac I. Genome-wide association mapping of milk production traits in Braunvieh cattle. Journal of Dairy Science 2012; 95: 5357–5364.

40. Lee SH, Choi BH, Lim D, Gondro C, Cho YM, Dang CG, et al. Genome-Wide Association Study Identifies Major Loci for Carcass Weight on BTA14 in Hanwoo (Korean Cattle). PLoS ONE 2013; 8: e74677.

41. Minozzi G, Nicolazzi EL, Stella A, Biffani S, Negrini R, Lazzari B, et al. Genome Wide Analysis of Fertility and Production Traits in Italian Holstein Cattle. PLoS ONE 2013; 8: e80219.

42. Yang J, Benyamin B, McEvoy BP, Gordon S, Henders AK, Nyholt DR, et al. Common SNPs explain a large proportion of the heritability for human height. Nat. Genet. 2010; 42: 565–569.

43. Yang J, Manolio TA, Pasquale LR, Boerwinkle E, Caporaso N, Cunningham JN, et al. Genome partitioning of genetic variation for complex traits using common SNPs. Nat. Genet. 2011; 43: 519–525.

44. Sutter NB, Bustamante CD, Chase K, Gray MM, Zhao K, Zhu L, et al. A Single IGF1 Allele Is a Major Determinant of Small Size in Dogs. Science 2007; 316: 112–115.

45. Oksenberg JR, Baranzini SE, Sawcer S, Hauser SL. The genetics of multiple sclerosis: SNPs to pathways to pathogenesis. Nat. Rev. Genet. 2008; 9: 516–526.

46. Thomas D. Gene—environment-wide association studies: emerging approaches. Nat. Rev. Genet. 2010; 11: 259–272.

47. Vilhjálmsson BJ, Nordborg M. The nature of confounding in genome-wide association studies. Nat. Rev. Genet. 2013; 14: 1–2.

48. Mackay TFC, Richards S, Stone EA, Barbadilla A, Ayroles JF, Zhu D, et al. The Drosophila melanogaster Genetic Reference Panel. Nature 2012; 482: 173–178.

49. Huang W, Massouras A, Inoue Y, Pfeiffer J, Ramia M, Tarone AM; et al. Natural variation in genome architecture among 205 Drosophila melanogaster Genetic Reference Panel lines. Genome Res.2014; 24: 1193–1208.

50. Ayroles JF, Carbone MA, Stone EA, Jordan KW, Lyman RF, Magwire MM, et al. Systems genetics of complex traits in Drosophila melanogaster. Nat. Genet. 2009: 41: 299–307.

51. Massouras A, Waszak SM, Albarca-Aguilera M, Hens K, Holcombe W, Ayroles JF, et al. Genomic Variation and Its Impact on Gene Expression in Drosophila melanogaster. PLoS Genet. 2012; 8: p. e1003055.

52. Nijhout HF, Riddiford LM, Mirth C, Shingleton AW, Suzuki Y, Callier V. The developmental control of size in insects. WIRES Dev. Biol. 2014; 3: 113–134.

53. Lippert C, Listgarten J, Liu Y, Kadie CM, Davidson RI, Heckerman D. FaST linear mixed models for genome-wide association studies. Nat. Methods 2011; 8: 833–835.

54. Bronstein R, Levkovitz L, Yosef N, Yanku M, Ruppin E, Sharan R, et al. Transcriptional regulation by CHIP/LDB complexes. PLoS Genet. 2010; 6: e1001063.

55. Franceschini A, Szklarczyk D, Frankild S, Kuhn M, Simonovic M, Roth A, et al. STRING v9.1: protein-protein interaction networks, with increased coverage and integration. Nucleic Acids Res. 2012; 41: D808–D815.

56. Liu JZ, Mcrae AF, Nyholt DR, Medland SE, Wray NR, Brown KM, et al. A Versatile Gene-Based Test for Genome-wide Association Studies. Am. J. Hum. Genet. 2010; 87: 139–145.

57. Schüpbach T, Xenarios I, Bergmann S, Kapur K. FastEpistasis: a high performance computing solution for quantitative trait epistasis. Bioinformatics 2010; 26: 1468–1469.

58. Müller P, Kuttenkeuler D, Gesellchen V, Zeidler MP, Boutros M. Identification of JAK/STAT signaling components by genome-wide RNA interference. Nature 2005; 436: 871–875.

59. Wang H, Chen X, He T, Zhou Y, Luo H. Evidence for tissue-specific Jak/STAT target genes in Drosophila optic lobe development. Genetics 2013; 195: 1291–1306.

60. Yang L, Meng F, Ma D, Xie W, Fang M. Bridging Decapentaplegic and Wingless signaling in Drosophila wings through repression of naked cuticle by Brinker. Development 2012; 140, 413–422.

61. Madan LL, Veeranna S, Shameer K, Reddy CCS, Sowdhamini R, Gopal B. Modulation of Catalytic Activity in Multi-Domain Protein Tyrosine Phosphatases. PLoS ONE 2011; 6: e24766.

62. Carter GW. Inferring gene function and network organization in Drosophila signaling by combined analysis of pleiotropy and epistasis. G3 2013; 3: 807–814.

63. Murali T, Pacifico S, Yu J, Guest S, Roberts GG 3rd, Finley RL Jr. DroID 2011: a comprehensive, integrated resource for protein, transcription factor, RNA and gene interactions for Drosophila. Nucleic Acids Res. 2011; 39: D736–D743.

64. Yu J, Pacifico S, Liu G, Finley RL Jr. DroID: the Drosophila Interactions Database, a comprehensive resource for annotated gene and protein interactions. BMC Genomics. 2008; 9: p. 461.

65. Celniker SE, Dillon LAL, Gerstein MB, Gunsalus KC, Henikoff S, Karpen GH, et al. Unlocking the secrets of the genome. Nature 2009; 459: 927–930.

66. Young RS, Marques AC, Tibbit C, Haerty W, Basset AR, Liu JL, et al. Identification and Properties of 1,119 Candidate LincRNA Loci in the Drosophila melanogaster Genome. Genome Biol. Evol. 2012; 4: 427–442.

67. Hangauer MJ, Vaughn IW, McManus MT. Pervasive Transcription of the Human Genome Produces Thousands of Previously Unidentified Long Intergenic Noncoding RNAs. PLoS Genetics 2013; 9: e1003569.

68. Ernst J, Kheradpour P, Mikkelsen TS, Shoresh N, Ward LD, Epstein CB, et al. Mapping and analysis of chromatin state dynamics in nine human cell types. Nature 2011; 473: 43–49.

69. Altschul SF, Gish W, Miller W, Myers EW, Lipman DJ. Basic local alignment search tool. J. Mol. Biol. 1990; 215: 403–410.

70. Hu Y, Flockhart I, Vinayagam A. Bergwitz C, Berger B, Perrimon N, et al. An integrative approach to ortholog prediction for disease-focused and other functional studies. BMC Bioinformatics 2011; 12: 357.

71. Durham MF, Magwire MM, Stone EA, Leips J. Genome-wide analysis in Drosophila reveals age-specific effects of SNPs on fitness traits. Nat. Commun. 2014; 5: p. 4338.

72. Carrington JC, Ambros V. Role of microRNAs in plant and animal development. Science 2003; 301: 336–338.

73. Inui M, Martello G, Piccolo S. MicroRNA control of signal transduction. Nat. Rev. Mol. Cell Biol. 2010; 11: 252–263.

74. Fatica A, Bozzoni I. Long non-coding RNAs: new players in cell differentiation and development. Nat. Rev. Genet. 2014; 15: 7–21.

75. Schleich S, Strassburger K, Janiesch PC, Koledachkina T, Miller KK, Haneke K et al. DENR-MCT1 promotes translation re-initiation downstream of uORFs to control tissue-growth. Nature 2014; 512: 208–212.

76. Schertel C, Huang D, Björklund M, Bischof J, Yin D, Li R, et al. Systematic Screening of a Drosophila ORF Library in vivo Uncovers Wnt/Wg Pathway Components. Dev. Cell 2013; 25: 207–219.

77. Unckless RL, Rottschaefer SM, Lazzaro BP. A Genome-Wide Association Study for Nutritional Indices in Drosophila. G3 2015; 5: 417–425.

78. Iida H, Nakamura H, Ono T, Okumura MS, Anraku Y. MID1, a novel Saccharomyces cerevisiae gene encoding a plasma membrane protein, is required for Ca2+ influx and mating. Mol Cell Biol 1994; 14: 8259–8271.

79. Levin DE, Errede B. The proliferation of MAP kinase signaling pathways in yeast. Curr. Opin. Cell Biol. 1995; 7: 197–202.

80. Syed ZA, Härd T, Uf A, van Dijk-Härd IF. A potential role for Drosophila mucins in development and physiology. PLoS ONE 2008; 3: e3041.

81. Povelones M, Howes R, Fish M, Nusse R, Genetic Evidence That Drosophila frizzled Controls Planar Cell Polarity and Armadillo Signaling by a Common Mechanism. Genetics 2005; 171: 1643–1654.

82. Parsons LM, Grzeschik NA, Allott M, Richardson H. Lgl/aPKC and Crb regulate the Salvador/Warts/Hippo pathway. Fly 2010; 4: 288–293.

83. Lin C, Katanaev VL. Kermit Interacts with Gao, Vang, and Motor Proteins in Drosophila Planar Cell Polarity. PLoS ONE 2013; 8: e76885.

84. Hatakeyama J, Wald JH, Printsev I, Ho HYH, Carraway KL. Vangl1 and Vangl2: planar cell polarity components with a developing role in cancer. Endocrine Related Cancer 2014; 21: R345–R356.

85. Weber U, Pataki C, Mihaly J, Mlodzik M. Combinatorial signaling by the Frizzled/PCP and Egfr pathways during planar cell polarity establishment in the Drosophila eye. Dev. Biol. 2008; 316: 110–123.

86. Sing A, Tsatskis Y, Fabian L, Hester I, Rosenfeld R, Serrichio M, et al. The Atypical Cadherin Fat Directly Regulates Mitochondrial Function and Metabolic State. Cell 2014; 158: 1293–1308.

87. van Bon BWM, Oortveld MAW, Nijtmans LG, Fenckova M, Nijhof B, Besseling J, et al. CEP89 is required for mitochondrial metabolism and neuronal function in man and fly. Hum. Mol. Genet. 2013; 22: 3138–3151.

88. Pereira C, Queirós S, Galaghar A, Sousa H, Pimentel-Nunes P, Brandão C, et al. Genetic variability in key genes in prostaglandin E2 pathway (COX-2, HPGD, ABCC4 and SLCO2A1) and their involvement in colorectal cancer development. PLoS ONE 2014; 9: e92000.

89. Clough E, Jimenez E, Kim YA, Whitworth C, Neville MC, Hempel LU, et al. Sex-and Tissue-Specific Functions of Drosophila Doublesex Transcription Factor Target Genes. Dev. Cell 2014; 31: 761–773.

90. Palenzona DL, Alicchio R. Differential response to selection on the two sexes in Drosophila melanogaster. Genetics 1973; 74: 533–542.

91. Menezes BF, Vigoder FM, Peixoto AA, Varaldi J. The influence of male wing shape on mating success in Drosophila melanogaster. Animal Behaviour 2013; 85: 1217–1223.

92. Houle D, Mezey J, Galpern P, Carter A. Automated measurement of Drosophila wings. BMC Evol. Biol. 2003; 3: p.25.

93. Turner SD. qqman: an R package for visualizing GWAS results using Q-Q and manhattan plots. biorXiv DOI: 10.1101/005165., http://cran.r-project.org/web/packages/qqman/index.html accessed 19.09.2014.

94. Dietzl G, Chen D, Schnorrer F, Su KC, Barinova Y, Fellner M et al. A genome-wide transgenic RNAi library for conditional gene inactivation in Drosophila. Nature 2007; 448: 151–156.

95. Krzywinski M, Schein J, Birol I, Connors J, Gascoyne R, Horsman D, et al. Circos: An information aesthetic for comparative genomics. Genome Res. 2009; 19: 1639–1645.

96. St Pierre SE, Ponting L, Stefancsik R, McQuilton P, FlyBase Consortium. FlyBase 102--advanced approaches to interrogating FlyBase. Nucleic Acids Res. 2014; 42: D780–8.

97. Clark AG, Eisen MB, Smith DR, Bergman CM, Oliver B, Markow TA, et al. Evolution of genes and genomes on the Drosophila phylogeny. Nature 2007; 450: 203–218.

